# Erdr1 orchestrates macrophage polarization and determines cell fate via dynamic interplay with YAP1 and Mid1

**DOI:** 10.1101/2023.09.17.557960

**Authors:** Yuhang Wang

## Abstract

Erythroid differentiation regulator 1 (Erdr1) is a stress-induced, widely distributed, extremely conserved secreted factor found in both humans and mice. Erdr1 is highly linked with the Hippo-YAP1 signaling. Initially identified as an inducer of hemoglobin synthesis, it has emerged as a multifunctional protein, especially in immune cells. Although Erdr1 has been implicated in T cells and NK cell function, its role in macrophage remains unclear. This study aims to explore the function and mechanism of Erdr1 in IL-1β production in macrophages. Data manifest Erdr1 could play an inhibition role in IL-1β production, which also has been reported by previous research. What significance is we discovered Erdr1 can promote IL-1β production which is associated with Erdr1 dose and cell density. We observed that Erdr1 was inhibited in pro-inflammatory (M1) macrophages but was upregulated in anti-inflammatory (M2) macrophages compared to naive macrophages. We hypothesized that Erdr1 dual drives and modulates IL-1β production by binding with distinct adaptors via concentration change. Mechanistically, we demonstrated that Erdr1 dual regulates IL-1β production by dynamic interaction with YAP1 and Mid1 by distinct domains. Erdr1-YAP1 interplay mediates macrophage M2 polarization by promoting an anti-inflammatory response, enhancing catabolic metabolism, and leading to sterile cell death. Whereas, Erdr1-Mid1 interplay mediates macrophage M1 polarization by initiating a pro-inflammatory response, facilitating anabolic metabolism, and causing inflammatory cell death. This study highlights Erdr1 orchestrates macrophage polarization and determines cell date by regulating YAP1 through non-classical Hippo pathway.

## Introduction

Erdr1 was initially identified as a stress-induced factor that acts as a hemoglobin synthesis-inducing factor in leukemia cells and was extremely conserved between humans and mice (1, 2). Subsequent research has demonstrated its important role in immune response (3–6). Erdr1 has been shown to play a critical role in T cell proliferation, activation (3, 4), and apoptosis (5). Recombinant Erdr1 protein treatment strengthens TCR signal in T cells by enhancing calcium influx (3) and PLCɣ1 signaling (4) upon activation by CD3 antibody *in vitro*, while *in vivo*, it significantly promotes thymocyte proliferation (3). However, erdr1 was also found to promote Treg cell activation (7), and endogenous overexpression of Erdr1 induces T cell apoptosis upon activation (5). The conflicting functions of Erdr1 on T cells are most likely due to the Erdr1 concentration and cell context. Apart from T cells, Erdr1 protein has also been shown to boost NK cell activation by promoting F-actin accumulation (6). Additionally, Erdr1 has been implicated in the pathogenesis of inflammatory disorders such as psoriasis (8) and rosacea (9). Notably, Erdr1 was downregulated in these patients and has a negative correlation with the pro-inflammatory cytokine IL-18 (9, 10), a readout cytokine of the inflammasome (11, 12). Animal experiments have manifested the broadly anti-inflammatory effect of Erdr1 protein in treating inflammatory disorders such as psoriasis (10), rosacea (9), colitis (13), and arthritis (7). Erdr1 has also been reported to play a negative role in macrophage migration(14). However, the function and mechanism of Erdr1 in macrophages are still unknown. Our preliminary data manifested that Erdr1 acts as an anti-inflammatory factor and could play an inhibition role in LPS-induced IL-1β production, which is consistent with previous reports. Interestingly, we discovered that Erdr1 also can facilitate IL-1β production. The dual role of Erdr1 over IL-1β production is correlated with Erdr1 dose and cell density. Particularly, we found that at decent cell density, Erdr1 is significantly down-regulated in LPS-induced pro-inflammatory macrophages (M1) but dramatically upregulated in IL-4-induced anti-inflammatory macrophages (M2) compared to naive macrophages (M0). The correlation of Erdr1 level with IL-1β production in macrophages follows a bell-shaped model. We hypothesize that, compared to basal Erdr1 level in steady state (M0), the up-regulated Erdr1 inhibits IL-1β production and mediates macrophage M2 polarization, while down-regulated Erdr1 promotes IL-1β production and mediates macrophage M1 polarization, which was achieved by interacts with different adaptors via Erdr1 conformation change under different concentration. We aim to investigate the candidates’ adaptors of Erdr1 and uncover the mechanism of Erdr1’s dual role in IL-1β production and macrophage M1/M2 polarization.

Previous research has demonstrated that secreted Erdr1 mediates cellular adaptation, it promotes cell survival at a low concentration and low cell density but enhances cell death at high concentrations and high cell density (1), indicating Erdr1 involved in maintaining cellular homeostasis. Furthermore, the observation that the Erdr1 C-terminal domain was undetectable at cell-cell contact sites suggested that it may be inactivated at these sites (1). Cell density-dependent inhibition, also known as contact inhibition, is a mechanism that regulates cell proliferation by modulating the Hippo-YAP1 pathway (15–17), which inspired us to link Erdr1 with the YAP1 signaling pathway. Moreover, through searching the database, we discovered the strong correlation of Erdr1 with YAP1 in various studies (5, 18–21). Firstly, Erdr1 has been found to directly regulate YAP1 signaling. For instance, in T cells, the overexpression of Erdr1 augments the expression of YAP1 and its signature gene, Amotl1, following T-cell stimulation(5). Secondly, YAP1 knockout has also been shown to impact Erdr1 expression. In Treg cells, YAP1 knockout prominently upregulates Erdr1 upon stimulation (19), and similarly, YAP1 knockout leads to a remarkable increase in Erdr1 expression in Müller glia cells during degeneration induction (20). However, it is worth noting that YAP1 knockout in mouse incisor epithelium results in a significant downregulation of Erdr1 (18). These intriguing findings suggest that YAP1 may regulate Erdr1 in a cell type-specific manner. Given the robust correlation of Erdr1 with YAP1, we hypothesized YAP1 is one of the binding adaptors of Erdr1. Cell density-dependent inhibition was also observed play role in controlling inflammation (22, 23), we hypothesized that Erdr1 regulates IL-1β production by directly modulating YAP1 in macrophages.

YAP1, the core of hippo pathway, has been established to mediate macrophage polarization and facilitate pro-inflammatory cytokine production in macrophages (24–26). The function of YAP1 is intricately linked to its expression levels and subcellular localization. YAP1 ON (Hippo OFF) occurs when YAP1 is translocated into the cell nuclear, where it interacts with its transcriptional co-activators, thereby facilitating the activation of the YAP1 signaling pathway. Conversely, YAP1 OFF (Hippo ON) achieved by capturing YAP1 in the cytoplasm, leading to the inhibition of the YAP1 signaling pathway (27). We hypothesized that Erdr1 regulates IL-1β production achieving by controlling YAP1 ON/OFF. Our preliminary data proved Erdr1 has a strong interaction with YAP1 in the cytoplasm in M2 but not M1 macrophages. We predicted Erdr1 can also drive YAP1 ON and promote M1 polarization by disconnecting with YAP1 and binding to another adaptor, Mid1. Mid1 is a highly conserved gene expressed ubiquitously in humans and mice (28). Initially identified for its essential role in development (28), Mid1 has also been shown to play a crucial role in macrophage activation (29–31). Our hypothesis is based on several lines of evidence. Firstly, Erdr1 and Mid1 are closely linked in gene mapping, as they are neighbor genes located in the PAR region of the sex chromosomes X and Y in mice (32–34). Secondly, Mid1 is functionally linked to Erdr1 (19, 35–52), YAP1 (19, 53, 54), and YAP1 downstream genes such as CTGF (55), Cry61 (35, 40, 44, 55), and IL-1β (31, 36, 55), as shown by previous microarray data and experiments. Thirdly, Erdr1 and Mid1 are known to play a shared role in regulating YAP1 signature gene expression (36, 55, 56). Fourthly, Mid1 is a zinc finger protein with three zinc finger domains and multiple zinc binding sites that are critical for its function. Erdr1 contains the HESTH sequence (118-122), representing a potent zinc-binding motif known as HEXXH (57). Both Erdr1 and Mid1 act as early stress sensors (1, 58) and involved in dysregulated intracellular zinc signaling (38) , and it is highly probable that they play a role in initial intracellular zinc mobilization through the formation of the Erdr1-Mid1 zinc channel. Overall, our hypothesis is that Erdr1 interacts dynamically with YAP1 and Mid1 to orchestrate macrophage activation and to determine the extent of inflammation. As an intrinsically disordered protein (IDP) (59), Erdr1 lacks of fixed structure and has the potential to play multiple functions by binding partners highly dynamically and cooperatively.

Mechanistically, we evidenced that Erdr1 modulates IL-1β production through dynamically interacting with YAP1 and Mid1 by distinct domains. Upregulated Erdr1 acts as an inhibitor of IL-1β production by sequestering YAP1 directly in the cytoplasm (YAP1 OFF), driving macrophage M2 polarization. Conversely, downregulated Erdr1 acts as an activator for IL-1β production via disconnecting with YAP1 but interacting with Mid1, ultimately driving YAP1 translocate to the nuclear (YAP1 ON), which promotes macrophage M1 polarization. More importantly, we also manifested that Erdr1, by concentration change, decides metabolism strength and mediates macrophage metabolic polarization towards a pro-anabolic or pro-catabolic state. Ultimately, Erdr1 decides cell fate, determining whether cells survive or undergo programmed cell death through either a pro-apoptotic or pro-pyroptotic pathway. This study highlights that Erdr1 dynamically drives and orchestrates macrophage polarization and determines cell fate by regulating YAP1 through the non-classical Hippo pathway.

## Materials and Methods

### Cell culture

293T and Hela cells were cultured with DMEM medium in the presence of 10% FBS, 100 IU/ml penicillin, and 100 μg/ml streptomycin. RAW246.7 were cultured with RPMI-1640 medium in the presence of 10% FBS, 100 IU/ml penicillin, 100 μg/ml streptomycin.

### Mouse BMDMs

C57BL/6 mice were killed, and femurs were removed. Bone marrow was harvested by flushing with BMDM medium (RPMI-1640 medium supplemented with 10% FBS, antibiotics, l-glutamine, and 10 μg/ml M-CSF). The harvested cells were centrifuged at 300 × g for 10 min and were treated with ACK lysis buffer (ThermoFisher, A1049201) for 1 min to remove RBCs. ACK lysis buffer was neutralized by adding 10 volumes of PBS, and cells were spun down at 300 × g for 10 min. The pelleted cells were resuspended in BMDM medium and were plated at 0.6*10^6^ cells/ml for culture. The medium was replaced on days 4, 5, and 6 of culture before cell use on day 7. In some experiments, BMDMs were treated with indicated concentrations of LPS, IL-4, or purified Erdr1 proteins. Some experiment seeding different cell density as indicated. All animal studies were approved by the University of Iowa Animal Care and Use Committee and meet stipulations of the Guide for the Care and Use of Laboratory Animals.

### Plasmid preparation and transduction

Full length Erdr1 (Erdr1-177) and Erdr1 deficiency C-terminal 32 amino acid (Erdr1-145) with C-terminal HA tags was synthesized (GenScript) and cloned into pcDNA3.1 plasmid using In-Fusion cloning (Clontech). YAP1 and Mid1 with C-terminal Flag were synthesized (GenScript) and cloned into pcDNA3.1 plasmid using In-Fusion cloning (Clontech). Plasmids were sequenced before use. For transient expression, plasmids were transfected into 293 T cell or Hela cell using Lipofectamine™ 3000 Transfection Reagent (Life Technologies) according to the manufacturer’s instructions. Lentiviral vectors: For gene knockout, we prepared following transfer plasmid. Erdr1, Mid1 and YAP1 Crispr/Cas9 knockout transfer plasmid were purchased from GenScript; For Erdr1 overexpression, full length Erdr1 (Erdr1-177) C-terminal HA tags were cloned into pHAGE vector. Lentivirus were produced by transfer lentiviral vector (transfer plasmid, psPAX2 and pMD2.G) into 293T cells following transfection protocol. Once produced, lentivirus was used for infection macrophage with different MOI.

### Cell viability assay

Cell viability assessed by MTT assay. MTT assay kit was purchased from Abcam (ab211091) and were performed according to the manufacturer’s protocol.

### Coimmunoprecipitation (Co-IP) assay

Cells were harvested with ice-cold PBS. Cell pellets were then lysed in immunoprecipitation buffer (IP lysis buffer: 25 mM Tris-HCl pH 7.4, 150 mM NaCl, 1 mM EDTA, 1% NP-40 and 5% glycerol, 1 mM DTT, and 1× protease/phosphatase inhibitors) for 30 min on ice and cleared by centrifugation at 13,000 × g for 15 min. The Erdr1-HA&YAP1-Flag or Erdr1-HA&Mid1-Flag was immunoprecipitated by Anti-FLAG® M2 Magnetic Beads for 2h, rotating end over end at 4°C. After the beads were pelleted, washed three times with IP lysis buffer, the immunocomplexes containing the Flag tagged interaction protein were then prior to elution with 2× SDS-sample buffer for 5 min at 95°C, followed by Immunoblotting analysis.

### Recombinant Erdr1

Generation of recombinant Erdr1 was performed by following protocol. Briefly, Erdr1-177 and Erdr1-145 with C-terminal His-tag was cloned into the pET28a vector. E. coli was transformed with this plasmid and cultured in medium containing ampicillin. IPTG was introduced for protein induction. Erdr1-177-His and Erdr1-145-His was purified on a nickel column and purity was >95% as confirmed by SDS-PAGE and Immunoblot. Potential endotoxin was removed using High-Capacity Endotoxin Removing Spin Columns (88274, ThermoFisher).

### Anti-Erdr1 polyclonal antibody

Anti-Erdr1 peptide Ab (36–51 amino acids of Erdr1, N-RAPRPPRHTRHTRHTR-C) was got as a gift from Dr. June Round at Utah University(5). Erdr1 neutralizing antibody were got as a gift from Dr. Timothy L. Denning (13).

### Immunofluorescence

Cells were seeded into 24 well plates containing 12 mm coated coverslips (Platinum Line). After indicated culture or treatment times, cells were fixed by incubation with PBS containing 4% (w/v) paraformaldehyde (Merck, #104005100) for 15 min at room temperature. After washing three times with PBS for 5 min, fixed cells were blocked and permeabilized by incubation with 500 μl methanol for 5 min. PBS containing 5% (w/v) horse serum (Sigma, #A7906) were used for blocking for 1 h at room temperature. Primary antibodies were diluted in 5% (w/v) horse serum/PBS and applied to coverslips and incubated at 4°C overnight. After three washes with PBS, coverslips were incubated with secondary antibody diluted in 5% (w/v) horse serum/PBS at room temperature for 1 h. Cells were washed three times with PBS. Cells were stained nucleic and mounted onto glass slides using Antifade Mounting Medium with DAPI (VECTASHIELD). Slides were viewed using Zeiss LSM 710 Confocal Microscope.

### Immunoblotting

Total cell extracts were lysed using radioimmunoprecipitation (RIPA) assay buffer, which was supplemented with Complete Mini Protease Inhibitor Cocktail and Phosphatase Inhibitor Cocktail from Roche. The lysates were put on ice for a period of 30 minutes and vortexed every 5 minutes. Following centrifugation at 15,000 rpm for 15 minutes at 4°C, the supernatants were collected. Proteins were resolved on a SDS polyacrylamide gel and transferred to a 0.45-μm PVDF membrane. For immunoblot analysis, the following primary antibodies were used at 1000-fold dilution: Anti-Flag (20543-1-AP); Anti-HA (51064-2-AP); Anti-β-actin (CST3700); Anti-IL-β (AF-401-NA); Anti-YAP1 (sc-101199); Anti-Mid1 (PA5-38524); Anti-p-S6K (CST9204); Anti-p-AMPK (CST2535); Anti-t-AMPK (CST2532); Anti-Caspase1 (AG-20B-0042-C100); Anti-cleaved-Caspase3 (CST9661); Anti-pro-caspase3 (CST9662); Anti-cleaved-Caspase8 (CST8592) ; Horseradish peroxidase (HRP)-conjugated secondary antibodies were used depending on the host species of the primary antibodies.

### Intracellular free Zinc detection

Detecting intracellular zinc by using a commercial zinc indicator (FluoZin-3, AM, Invitrogen™ F24195). In brief, RAW246.7 cell was cultured in 96 wells plate with 0.6*10^6^ cell/ml and cultured overnight. Washed cell with PBS twice and then treated with 1 μg/ml LPS, 10 μM ZnCl_2_, indicated Erdr1 protein, and 1 μM FluoZin-3-AM in serum-free medium. Measure the fluorescence with (Ex/Em, 494/516) at indicated time points with well plate reader.

### Cytokine Quantification

IL-1β quantification was performed by harvesting cell supernatants at the indicated time points and treatment and assaying for IL-1β levels using a mouse IL-1β ELISA kit (DY401-050), according to the manufacturer’s protocol.

### qPCR and primers

Total RNA was extracted from macrophages at the indicated time points and treatment using TRIzol reagent (Invitrogen). 2 μg of RNA were used for cDNA synthesis. 2 μl of cDNA were added to 23 μl of PCR mixture containing 2xSYBR Green Master Mix (ABI) and 0.2 μM of forward and reverse primers. Amplification was performed in an ABI Prism 7500 thermocycler. Genes and forward (F) and reverse (R) primers are shown: IL-1β F: ACTGTTTCTAATGCCTTCCC; IL-1β R: TGGTTTCTTGTGACCCTGA; IL-18 F: GACTCTTGCGTCAACTTCAAGG; IL-18 R: CAGGCTGTCTTTTGTCAACGA; IL-6 F: TAGTCCTTCCTACCCCAATTTCC; IL-6 R: TTGGTCCTTAGCCACTCCTTC; TNF F: GACCCCTTTACTCTGACCCC; TNF R: AGGCTCCAGTGAATTCGGAA; IL-10 F: CGGGAAGACAATAACTGCACCC; IL-10 R: CGGTTAGCAGTATGTTGTCCAGC; TGF-β F: TGATACGCCTGAGTGGCTGTCT; TGF-β R: CACAAGAGCAGTGAGCGCTGAA; CTGF F: TGCGAAGCTGACCTGGAGGAAA; CTGF R: CCGCAGAACTTAGCCCTGTATG; Cyr61 F: GTGAAGTGCGTCCTTGTGGACA; Cyr61 R: CTTGACACTGGAGCATCCTGCA; Erdr1 F: TGATGTCACCCACGAAAGCA; Erdr1 R: TTCCTCCGTGAGAATCGCTC; Mid1 F: TGGACCTCAGAGGACGAGTT; Mid1 R: CCACGTTGACAACATTGGCT; YAP1 F: CGGCAGTCCTCCTTTGAGAT; YAP1 R: TTCAGTTGCGAAAGCATGGC; β-actin F: GTGGGAATGGGTCAGAAGGA; β-actin R: CTTCTCCATGTCGTCCCAGT. Cycle threshold (Ct) values were normalized to those of the housekeeping gene β-actin by the following equation: ΔCt = Ct (gene of interest) -Ct (β-actin). All results are shown as a ratio to β-actin calculated as 2−(ΔCt).

### Statistics

The p values of dynamic intracellular free zinc mobilization were determined by two-way ANOVA as specified in the figure legend. A Student’s t test was used to analyze differences in mean values between groups. All results are expressed as means ± SD. P values less than 0.05 were considered statistically significant; *P < 0.05; **P < 0.01; ***P < 0.001, ****P < 0.0001.

## Results

### Erdr1 is distinctly regulated in M1/M2 macrophages

Macrophages can be polarized into pro-inflammatory M1 macrophages and anti-inflammatory M2 macrophages (60, 61), which are the two extremes of a spectrum of macrophage phenotypes. We first investigated if Erdr1 expression was regulated during the macrophage polarization process. Here, we induced the polarization of M0 macrophages (Bone marrow-derived macrophage, BMDM) into M1 macrophages by stimulating with 1 μg/ml of LPS for 24 hours, or into M2 macrophages by stimulating with 20 ng/ml of IL-4 for 24 hours and confirming M1 marker genes (IL-1β, IL-6 and TNF) and M2 marker gene (IL10 and TGFβ) expression by qPCR (data not shown). We observed Erdr1 expression was dramatically reduced in M1 macrophages at mRNA level (Fig. 1A) and protein level (Fig. 1B), whereas it was significantly elevated in M2 macrophages (Fig. 1A and Fig. 1B), compared to M0 macrophages. Further investigation was conducted to assess the secretion and accumulation of Erdr1 in the supernatant of M1 and M2 macrophages, considering it was reported to secrete upon cellular stress. Immunoblot analysis revealed a dramatically elevated presence of Erdr1 in the supernatant of M2 macrophages when compared to M0 macrophages (Fig. 1B). Surprisingly, there was also a significantly increased level of Erdr1 detected in the supernatant of M1 macrophages (Fig. 1B), despite the decrease in endogenous Erdr1 expression (Fig. 1A and Fig. 1B). This increased secretion of Erdr1 in the medium of M1 macrophages might serve as a strategy to rapidly decrease the intracellular levels of Erdr1, rather than an induced expression factor. We conducted further investigations into the subcellular localization of Erdr1 in the polarized macrophage, employing immunofluorescence techniques. Our findings revealed Erdr1 has increased nuclear localization in M1 macrophage but increased cytoplasm localization in M2 macrophage compared to M0 macrophage (Fig. 1C). We further confirmed these observations by ImageJ analysis, with significantly augmented Erdr1 Nuclear/Cytoplasm intensity ratio in M1 macrophages, while a decreased ratio in M2 macrophages, compared to M0 (Fig. 1D). These findings manifest Erdr1 exhibits significant but distinct expression patterns and subcellular localization patterns in M1/M2 macrophage, implying Erdr1 might play roles both in M1 and M2 macrophages.

**Figure 1.**
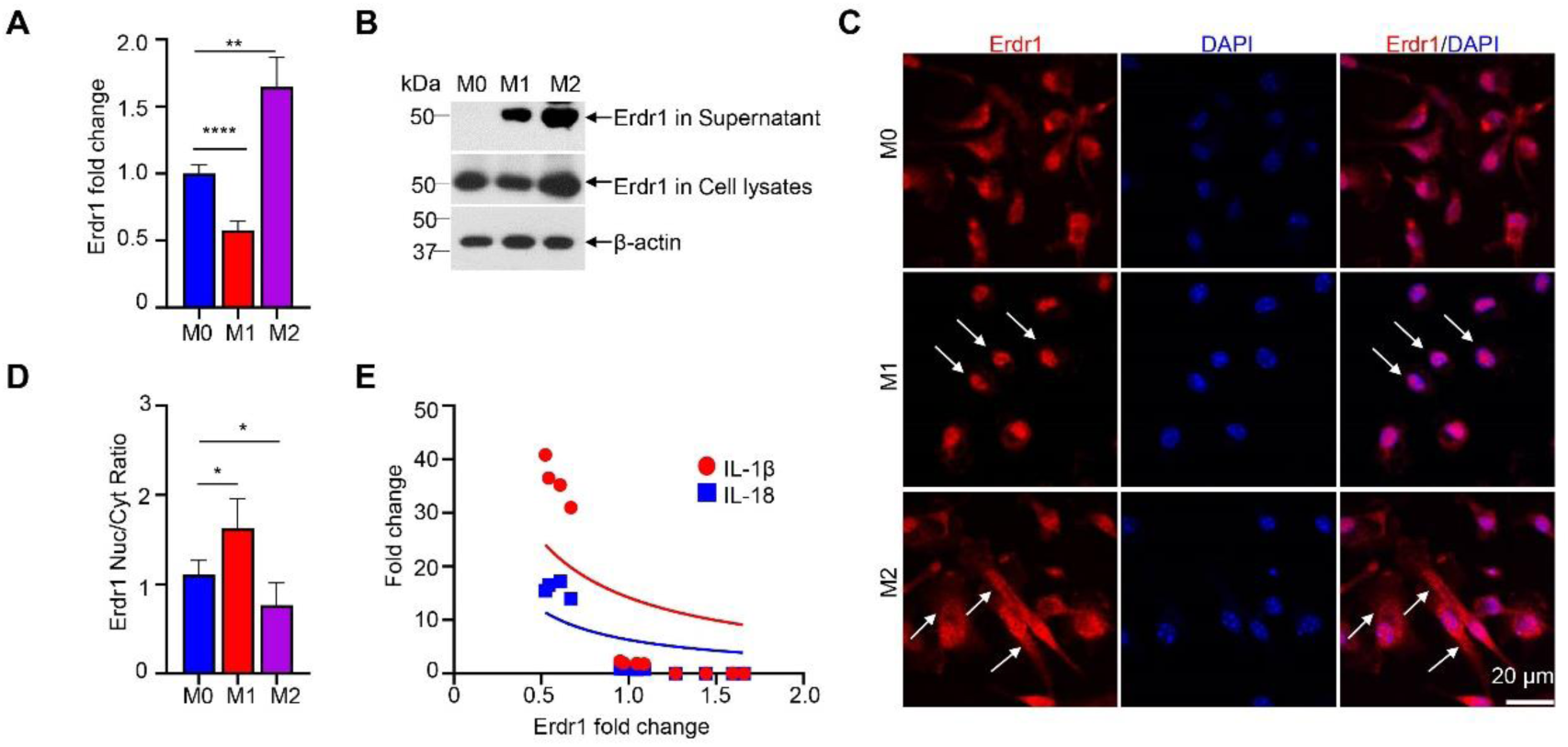
Erdr1 is distinctly regulated in M1/M2 macrophages. M0 macrophage used in this study was Bone marrow-derived macrophage (BMDM). Cell was seeded with density 0.6*10^6^ cells/ml in 24 well plates. M0 macrophage was induced into M1 macrophage by treating with 1 μg/ml LPS for 24 hours, or into M2 macrophage by treating with 20 ng/ml IL-4 for 24 hours. The same experiments were also conducted and got repeatable results in RAW246.7 cell. (**A)** qPCR analysis of Erdr1 mRNA expression in M0, M1, and M2 macrophages, qPCR normalized to β-actin expression (n=4). (**B)** Immunoblot analysis of Erdr1 level in cell lysates and supernatant of M0, M1, and M2 macrophages. β-actin as the internal control. (**C)** The Immunofluorescence staining of Erdr1 in M0, M1, and M2 macrophage and counterstained by DAPI. The white arrows in the M1 macrophage display Erdr1 is mainly localized in nuclear. The white arrows in the M2 macrophage display Erdr1 is mainly localized in cytoplasm. Scale bar in represents 20 μm. (**D)** Erdr1 Nuclear to Cytoplasm (Neu/Cyt) intensity ratio was quantified by ImageJ based on immunostaining. n=5 with more than 10 macrophages were analyzed per sample. (**E)** Curve Fitting correlation of Erdr1&IL-1β fold change and Erdr1&IL-18-fold change using Nonlinear Regression (n=4). Data represent mean ± SD. *p<0.05, **p<0.01, ****p<0.001. Data are representative of at least three independent experiments.

IL-1β is one of the most important pro-inflammatory cytokines induced in M1 macrophages (60, 61). IL-1β is processed during the Inflammasome activation process, as well as IL-18 (11, 12). Erdr1 has been shown to have a negative correlation with IL-18 (62). Consistent with that, here we demonstrated a negative correlation of endogenous Erdr1 level with IL-1β and IL-18 level in macrophages (Fig. 1E). IL-10, which is the anti-inflammatory cytokines induced in M2 macrophages, has positive correlation with Erdr1 (data not shown). This indicated Erdr1 is a signature gene for M2 macrophages, while if and how the downregulated Erdr1 plays role in M1 macrophages still not clear.

### Erdr1 has dose-dependent bell-shaped effect in LPS-induced IL-1β production in macrophages

We further aimed to elucidate the function of Erdr1 in regulating IL-1β production in M1 macrophages. Because we observed Erdr1 exhibits both endogenous regulation and induced secretion during macrophage activation, we conducted experiment to investigate how endogenous and exogenous Erdr1 affect the production of IL-1β in response to LPS stimulation.

First, we assessed the endogenous Erdr1 function on LPS-induced IL-1β production. M0 macrophages were genetically modified to either knockout or overexpress Erdr1 through transduction with Erdr1 CRISPR/Cas9 knockout or Erdr1 overexpression lentiviruses, respectively confirmed Erdr1 expression by qPCR (Fig. 2A and Fig. 2D). We observed that Erdr1 knockdown significantly suppressed LPS-induced IL-1β production at mRNA level in cell lysate (Fig. 2B) and protein level in the supernatant (Fig. 2C). Interestingly, we observed that moderate overexpression of Erdr1 (Erdr1 OE1, about 4.3 folds higher) dramatically enhanced LPS-induced IL-1β production, while a high-level overexpression of Erdr1 (Erdr1 OE2, about 28.6 folds higher) inhibited LPS-induced IL-1β production (Fig. 2E and Fig. 2F). Notably, we discovered a bell-shaped correlation between the endogenous Erdr1 and IL-1β production (Fig. 2G), suggesting that Erdr1 plays a dual role in regulating IL-1β production.

**Figure 2.**
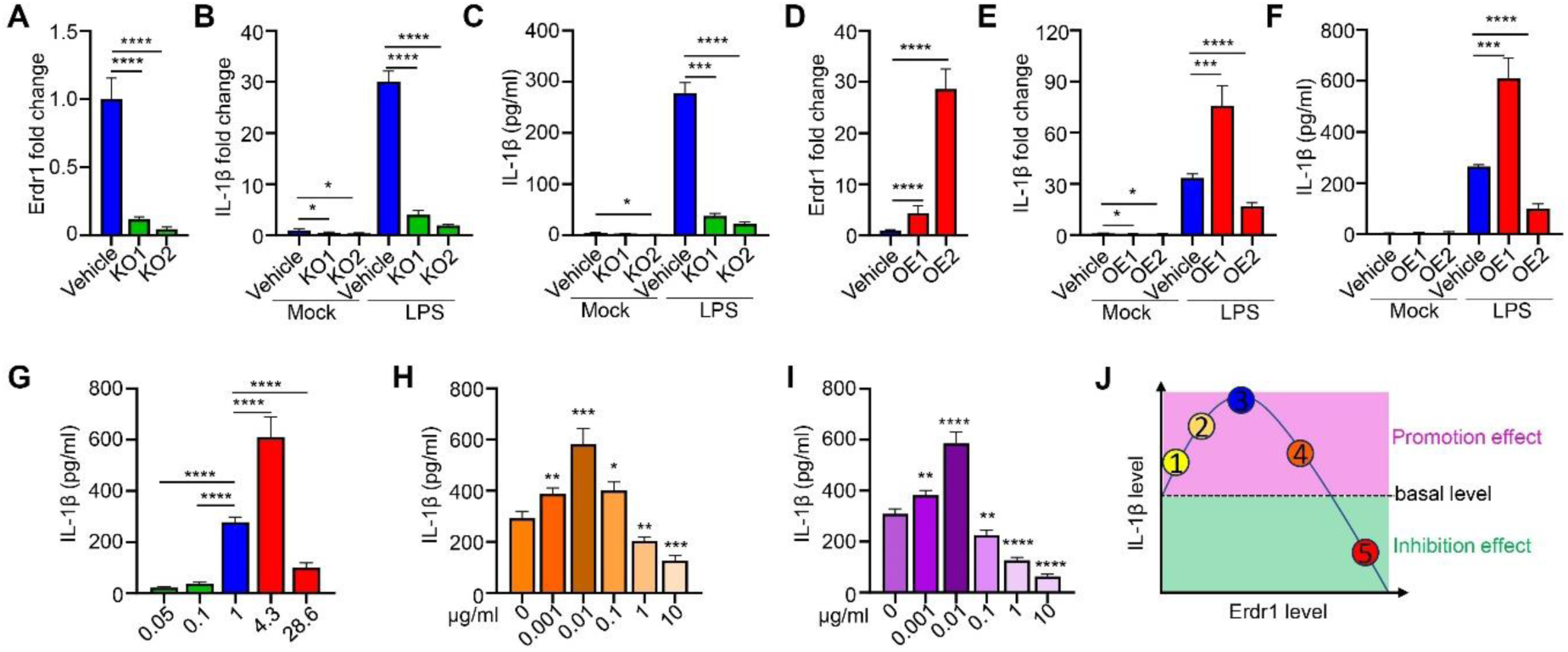
Erdr1 has bell-shape effect in LPS induced IL-1β production in macrophages. **(A-C)** Knockout of Erdr1 in BMDM by Lentivirus transduction for 40 hours, and then treated without or with 1 μg/ml LPS for 24 hours for the following analysis. Erdr1 KO1 and Erdr1 KO2 represent two knockout vectors, the vehicle is transfected with the control Lentivirus vector (n=4). **(A)** qPCR detecting Erdr1 expression after lentivirus transduction for 40 hours. (**B)** qPCR detecting IL-1β expression. (**C)** Detecting IL-1β production at supernatant by ELISA. (**D-F)** Overexpression of Erdr1 in BMDM by Lentivirus transduction for 40 hours and then treated without or with 1 μg/ml LPS for 24 hours for the following analysis. Erdr1 OE1 and Erdr1 OE2 represent two transfection titers of Lentivirus, the vehicle is transfected with the control Lentivirus vector (n=4). (**D)** qPCR detecting Erdr1 expression after lentivirus transduction for 40 hours. (**E)** qPCR detecting IL-1β expression. (**F)** Detecting IL-1β production at supernatant by ELISA. **(G)** The correlation of endogenous Erdr1 level (qPCR) and IL-1β level (ELISA) summary from (**C**) and (**F**). (**H)** Detecting IL-1β level at supernatant by ELISA in 1 μg/ml LPS-induced BMDM with the indicated concentration of Erdr1 antibodies treatment (n=4). (**I)** Detecting IL-1β level at supernatant by ELISA in 1 μg/ml LPS-induced BMDM with the indicated concentration of purified full-length Erdr1 protein treatment (n=4). (**J)** Threshold Model of Erdr1 in IL-1β Production: A Bell-Shaped Function in IL-1β Production. 1,2, low level of Erdr1 has positive correlation with IL-1β. 3, The optimal Erdr1 concentration promotes the peak expression of IL-1β. 4,5, high level of Erdr1 has negative correlation with IL-1β level. These show Erdr1 has both positive and negative roles in IL-1β production by dose change. (**A-I),** Cell density is 0.6*10^6^/ml and seeded in 24 well plates. All qPCR normalized to β-actin expression. Data represent mean ± SD. *p<0.05, **p<0.01, ***p<0.001 and, ****p<0.0001. Data are representative of three independent experiments.

Subsequently, we investigated the impact of the secreted Erdr1 in the medium on LPS-induced IL-1β production. To achieve this, we employed a range of concentrations of Erdr1 neutralization antibody to block the secreted Erdr1 in the medium. Interestingly, we observed a bell-shaped relationship between the Erdr1 antibody concentration and IL-1β production (Fig. 2H). These results suggest that an optimal concentration of Erdr1 in the medium exerts a positive effect on promoting IL-1β production. However, an excessive accumulation of Erdr1 in the medium appears to trigger a negative feedback mechanism, ultimately inhibiting IL-1β production.

Since purified Erdr1 protein has been shown to play an inhibition role in inflammation (8, 9, 63). We next tested Erdr1 protein effect in IL-1β production in macrophages through an exogenous protein supplementation experiment. Purified full-length Erdr1 was prepared and supplemented with a series of concentrations as indicated to the medium. Notably, data manifested Erdr1 protein has a dose-dependent bell-shaped effect in regulating LPS-induced IL-1β production (Fig. 2I). These findings provide striking evidence that a low dose of Erdr1 in the medium could induce IL-1β production, whereas a high dose of Erdr1 exerts an inhibitory role in IL-1β production. This indicates that the function of Erdr1 in IL-1β production is highly correlated to Erdr1 dose.

In summary, our data support that Erdr1 could play an inhibition role in IL-1β production, which has been reported. But we for the first time, demonstrate that Erdr1 also has the potential to promote IL-1β production. These data suggest the influence of Erdr1 in IL-1β production following a dose-dependent bell-shaped model (Fig. 2J).

### Erdr1 effect in IL-1β production also correlates with cell density in macrophages

Our observations above were obtained from samples with cell density of 0.6*10^6^ cells/ml. Given that prior research has reported Erdr1 effect in cell viability was influenced by cell density(1), we then investigated if Erdr1’s role in IL-1β expression was affected by cell density, and if Erdr1 affects cell viability in macrophages.

We seeded macrophages with 3 different densities, low density (0.1*10^6^ cells/ml), moderate density (0.5*10^6^ cells/ml), and high density (2.5*10^6^ cells/ml). Consistent with previous research (22, 23), we confirmed that both basal and LPS-stimulated IL-1β expression exhibited an increase at low density while being suppressed at higher density compared to levels at moderate density (Fig. 3A). We also observed that high cell density protects the cell from death (Fig. 3B). Interestingly, we noted cell density affects basal endogenous Erdr1 expression levels (Fig. 3C), Specifically, heightened basal Erdr1 expression was evident in scenarios of high cell density, whereas low cell density exhibited reduced Erdr1 expression, compared to the moderate one (Fig. 3C). Furthermore, we observed Erdr1 is upregulated under conditions of low cell density upon LPS stimulation, this response is contrary to our observations in both moderate and high-density scenarios (Fig. 3C). Additionally, we also noted that cell density influences the subcellular localization of Erdr1, it tends to localize in the cytoplasm at high density, while favors nuclear localization at low density (Supplemental Fig.1A and 1B). These findings reveal that basal and LPS-induced endogenous Erdr1 expression and subcellular localization were affected by cell density. Moreover, endogenous Erdr1 demonstrates a positive correlation with IL-1β at low cell density, while manifests a negative correlation with IL-1β at high cell densities upon LPS stimulation.

**Figure 3.**
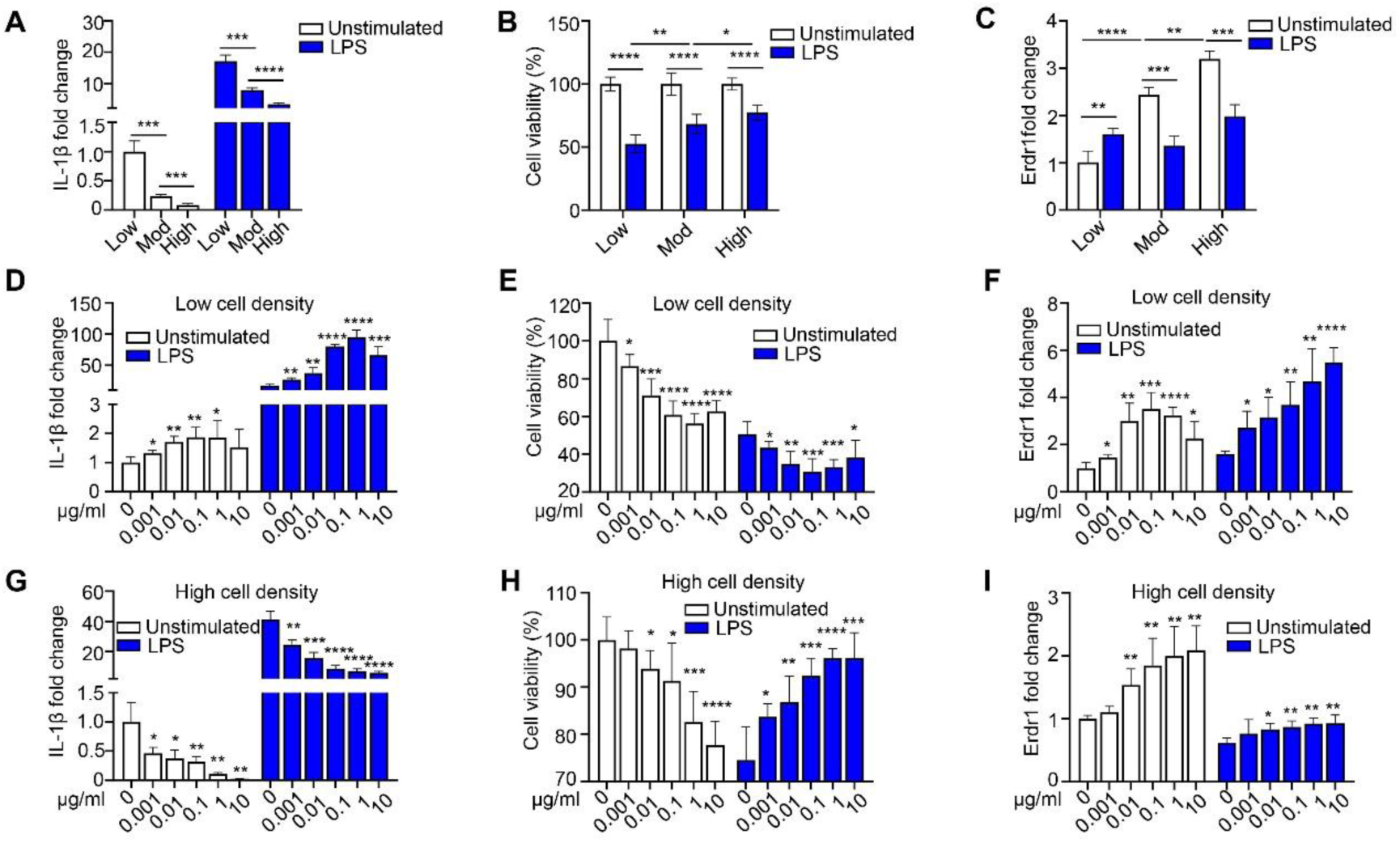
Erdr1 effect in IL-1β production correlates with cell density in macrophages. **(A-C)** BMDM were seeded with the indicated cell density: Low density (Low, 0.1*10^6^ cells/ml), Moderate density (Mod, 0.5*10^6^ cells/ml), or High density (High, 2.5*10^6^ cells/ml) in 24 well plates for 8 hours, and then treated without or with 1 μg/ml LPS for 24 hours or 48 hours (only for MTT assay) and perform following analysis: **(A)** Detecting IL-1β expression by qPCR (n=4). (**B)** Detecting cell viability by MTT assay (n=6). **(C)** Detecting Erdr1 expression by qPCR (n=4). (**D-F)** BMDM were seeded with Low cell density (0.1*10^6^ cells/ml) in 24 well plates for 8 hours, and then treated without or with 1 μg/ml LPS for 24 hours together with supplemented with the indicated concentration of Erdr1 (0, 0.001, 0.01, 0.1, 1, 10 μg/ml), and perform following analysis: (**D)** Detecting IL-1β expression by qPCR (n=4). **(E)** Detecting cell viability by MTT assay (n=6). **(F)** Detecting Erdr1 expression by qPCR (n=4). **(G-I)** BMDM were seeded with High cell density (2.5*10^6^ cells/ml) in 24 well plates for 8 hours, and then treated without or with 1 μg/ml LPS for 24 hours together with supplemented with the indicated concentration of Erdr1 (0, 0.001, 0.01, 0.1, 1, 10 μg/ml), and perform following analysis: **(G)** Detecting IL-1β expression by qPCR (n=4). **(H)** Detecting cell viability by MTT assay (n=6). **(I)** Detecting Erdr1 expression by qPCR (n=4). All qPCR normalized to β-actin expression. The same experiments were also conducted and got repeatable results in RAW246.7 cell. Data represent mean ± SD. *p<0.05, **p<0.01, ***p<0.001 and, ****p<0.0001. Data are representative of two independent experiments.

Subsequently, we investigated the impact of supplementing Erdr1 protein on both IL-1β expression and cell viability under conditions of low and high cell densities. The data clearly reveal that Erdr1 protein exerts a dose-dependent enhancing effect on IL-1β expression (Fig. 3D) and facilitates cell death exclusively at low cell density (Fig. 3E). In contrast, at high cell density, Erdr1 displays a dose-dependent inhibitory role in IL-1β expression (Fig. 3G), while also promotes cell survive after LPS stimulation (Fig. 3H). These observations strongly suggest that the function of Erdr1 in IL-1β modulation is profoundly influenced by cell density. Overall, Erdr1 emerges as an amplifier of pro-inflammatory signaling under conditions of low cell density yet transits into an inhibitor of pro-inflammatory signaling when cell density is high.

We proceeded to investigate the effects of Erdr1 protein on endogenous Erdr1 expression. Intriguingly, our data reveal that Erdr1 protein treatment led to an upregulation of endogenous Erdr1 in both low and high cell density settings (Fig. 3F and 3I). Notably, upon LPS stimulation, endogenous Erdr1 exhibits an upregulation in low cell density scenarios, a response further augments by Erdr1 protein treatment (Fig. 3F), ultimately intensifies IL-1β expression (Fig. 3D) and aggravates cell death (Fig. 3E). In contrast, under conditions of high cell density, LPS stimulation resulted in the downregulation of Erdr1, an effect that was counteracted by Erdr1 protein treatment (Fig. 3I), subsequently inhibits IL-1β expression (Fig. 3G) and reduces cell death (Fig. 3H). Remarkably, even in the absence of LPS stimulation, Erdr1 exhibits the ability to upregulate endogenous Erdr1 expression (Fig. 3F and 3I) and drive cell death (Fig. 3E and 3H) both at low and high cell density. These observations suggest endogenous Erdr1 plays a crucial role in maintaining cellular homeostasis. It becomes apparent that controlled fluctuations of endogenous Erdr1 level facilitate cellular survival. However, when Erdr1 levels veer to extremes-either too low or excessively high-it becomes a mediator of cell death.

In summary, these data clearly demonstrate that the effect of Erdr1 on IL-1β expression is closely correlated with cell density. And exogenous Erdr1 can manipulate IL-1β by regulates intricate Erdr1 level.

### Erdr1 promotes IL-1β production through activating YAP1 and Mid1 signaling pathway

We further to explore the mechanism of unique promotion role of Erdr1 in regulating IL-1β production. We hypothesized YAP1 and Mid1 are involved in this process as stated before.

To validate this hypothesis, we initially examined if and how YAP1 and Mid1 are regulated in M1/M2 macrophages. Notably, we observed remarkable upregulation of Mid1 in mRNA level (Fig. 4A) and protein level (Fig. 4B) in M1 macrophages compare to M0 macrophages. We also detected increased YAP1 expression in M1 macrophages at the protein level (Fig. 4B). In contrast, we observed a significant decreased expression of Mid1 (Fig. 4A and Fig. 4B) and YAP1 (Fig. 4B) in M2 macrophages. These findings indicate that Mid1 and YAP1 are also distinct regulated in M1/M2 macrophages. Utilizing immunofluorescence, we investigated the subcellular localization of YAP1 and Mid1 in M1/M2 macrophages. YAP1 has increased nuclear localization in M1 macrophages, but increased cytoplasm localization in M2 macrophages (Fig. 4C). Whereas we observed remarkable increased cytoplasm localization of Mid1 in M1 macrophages, but increased nuclear localization in M2 macrophages, compared to M0 macrophages (Fig. 4D). Further validated the observation through the quantification of the Nuclear to Cytoplasm intensity ratio (Nuc/Cyt ratio) of YAP1 and Mid1 using ImageJ. (Fig. 4F). These findings reveal that Mid1 and YAP1 also have dynamic regulation and distinct subcellular localization patterns during macrophage polarization.

**Figure 4.**
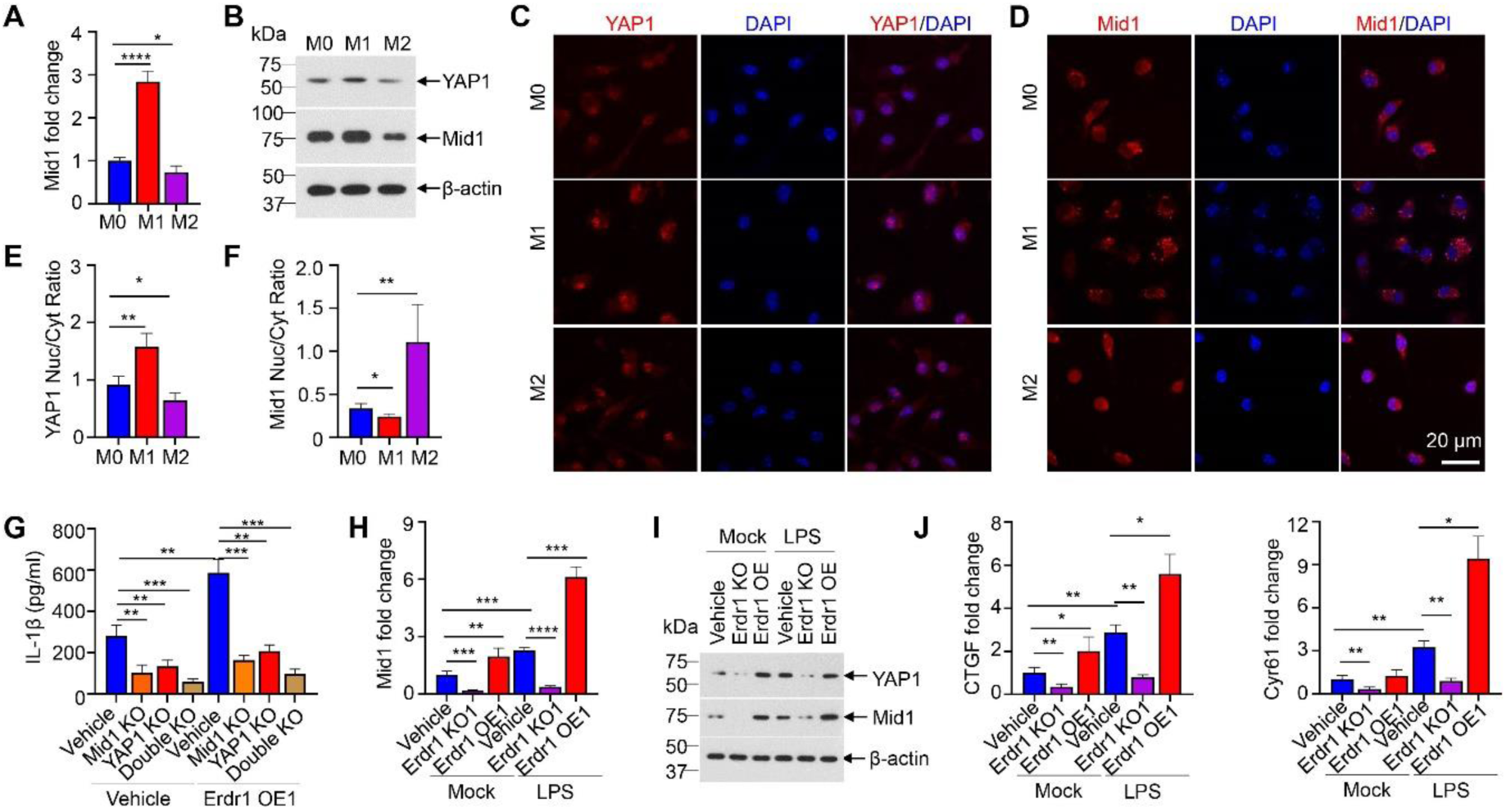
Erdr1 modulates IL-1β production through regulation of YAP1 and Mid1 signaling pathways. **(A)** qPCR analysis of Mid1 expression in M0, M1, and M2 macrophage (n=4). **(B)** Immunoblot analysis of YAP1 and Mid1 levels in the cell lysates of M0, M1, and M2 macrophage. **(C-D)** The Immunofluorescence staining of YAP1 (**C**) and Mid1 (**D**) in M0, M1, and M2 macrophage and counterstained by DAPI. Scale bar in represents 20 μm. **(E-F)** Nuclear to Cytoplasm (Neu/Cyt) intensity ratios YAP1 (**E**) and Mid1 (**F**) were quantified by ImageJ. n=5 with more than 10 macrophages were analyzed per sample. **(G)** Detecting IL-1β production by ELISA in indicated BMDM group before and after 1 μg/ml LPS stimulation for 24 hours (n=4). Mid1 KO, YAP1 KO, and Double KO represent knockout Mid1, YAP1, and both Mid1/YAP1 in BMDM by transfection CRISPR/cas9 Lentivirus; Erdr1 OE1 represent moderate overexpression of Erdr1 (about 4 folds) in BMDM by transfection Lentivirus; the vehicle is transfected with the control Lentivirus vector (n=4). (**H-J)** BMDM were knockout Erdr1 (Erdr1 KO1) or moderate overexpressed Erdr1 (Erdr1 OE1), then treated without or with 1 μg/ml LPS for 24 hours for following analysis. (**H)** qPCR analysis of Mid1 expression(n=4); **(I)** Immunoblot analysis of YAP1 and Mid1 expression. **(J)** qPCR analysis of CTGF (left) and Cyr61 (right) (n=4). All qPCR normalized to β-actin expression. (**A-J**) Cell density is 0.6*10^6^/ml and seeded in 24 well plates. Data represent mean ± SD. *p<0.05, **p<0.01, ***p<0.001 and, ****p<0.0001. Data are representative of two or three independent experiments.

We further investigated if YAP1 and Mid1 are involved in Erdr1-mediated LPS-induced IL-1β production. Data demonstrate that knockout of Mid1 or YAP1 or both led to significant blockade of IL-1β production in macrophages induced by LPS (Fig. 4F), indicating Mid1 and YAP1 are both essential for IL-1β production in M1 macrophages. Moderate overexpression of Erdr1 (Erdr1 OE1) has been demonstrated promote LPS-induced IL-1β production in macrophage (Fig. 2F), however, this promotion role is effectively blocked in Mid1 knockout, YAP1 knockout, or Mid1/YAP1 double knockout macrophages (Fig. 4G). These suggest Erdr1 plays roles in promoting IL-1β production by YAP1 and Mid1 signaling pathways.

We next tested if Erdr1 is the upstream regulator of the YAP1 and Mid1 signaling. Our data display that Erdr1 OE1 macrophage which enhances LPS-induced IL-1β production (Fig. 2F), also promotes LPS-induced Mid1 and YAP1 expression (Fig. 4H and Fig. 4I). Whereas, in Erdr1 deficiency macrophage, which blocks IL-1β production, Mid1 and YAP1 expression are both dramatic suppressed (Fig. 4H and Fig. 4I). These data manifest that Erdr1 regulates IL-1β production is achieved by direct regulating YAP1 and Mid1 expression. Furthermore, we prove other YAP1 and Mid1 downstream genes, such as CTGF and Cyr61 (55, 64, 65), are also regulated by Erdr1 (Fig. 4J), with Erdr1 overexpression (Erdr1 OE1) promotes CTGF and Cyr61 expression, but they are inhibited in Erdr1 deficiency macrophage (Fig. 4J), providing additional evidence that Erdr1 acts as an upstream regulator of YAP1 and Mid1.

Taken together, these findings provide compelling evidence that Erdr1 plays a crucial role in promoting IL-1β production by activating YAP1 and Mid1 signaling pathways in macrophages.

### Erdr1 displays distinct interaction with YAP1 and Mid1 by different domains and mediates macrophage polarization

We hypothesized Erdr1 plays dual role in IL-1β production by distinct and dynamic interaction with YAP1 and Mid1. To investigate this, we employed Co-immunoprecipitation (Co-IP) to detect the potential interaction of Erdr1 with YAP1 and Mid1 by using 293T cells. Specifically, we hypothesized that the C-terminal domain of Erdr1 is the region responsible for interacting with YAP1. This hypothesis is grounded in previous research, which has shown that the Erdr1 C-terminal 81 amino acids are involved in contact inhibition (2). Though protein sequence and function prediction via UniProt, we predicted the C-terminal 32 amino acids are critical for YAP1 binding. We observed full-length Erdr1 (Erdr1-177) has direct interaction with YAP1, whereas the Erdr1 variant which lacking the C-terminal 32 amino acids (Erdr1Δ32, or Erdr1-145) (Fig 6A) fails to exhibit this interaction (Fig. 5A). These reveal Erdr1 has direct interaction with YAP1 by its C-terminal 32 amino acid. Furthermore, our data manifest Mid1 could direct interaction with Erdr1-177 and even a stronger interaction with Erdr1-145 in Co-IP experiments (Fig. 5B), suggesting Erdr1 has the potential to interact with Mid1, particularly when it exposes its Erdr1-145 domain.

**Figure 5.**
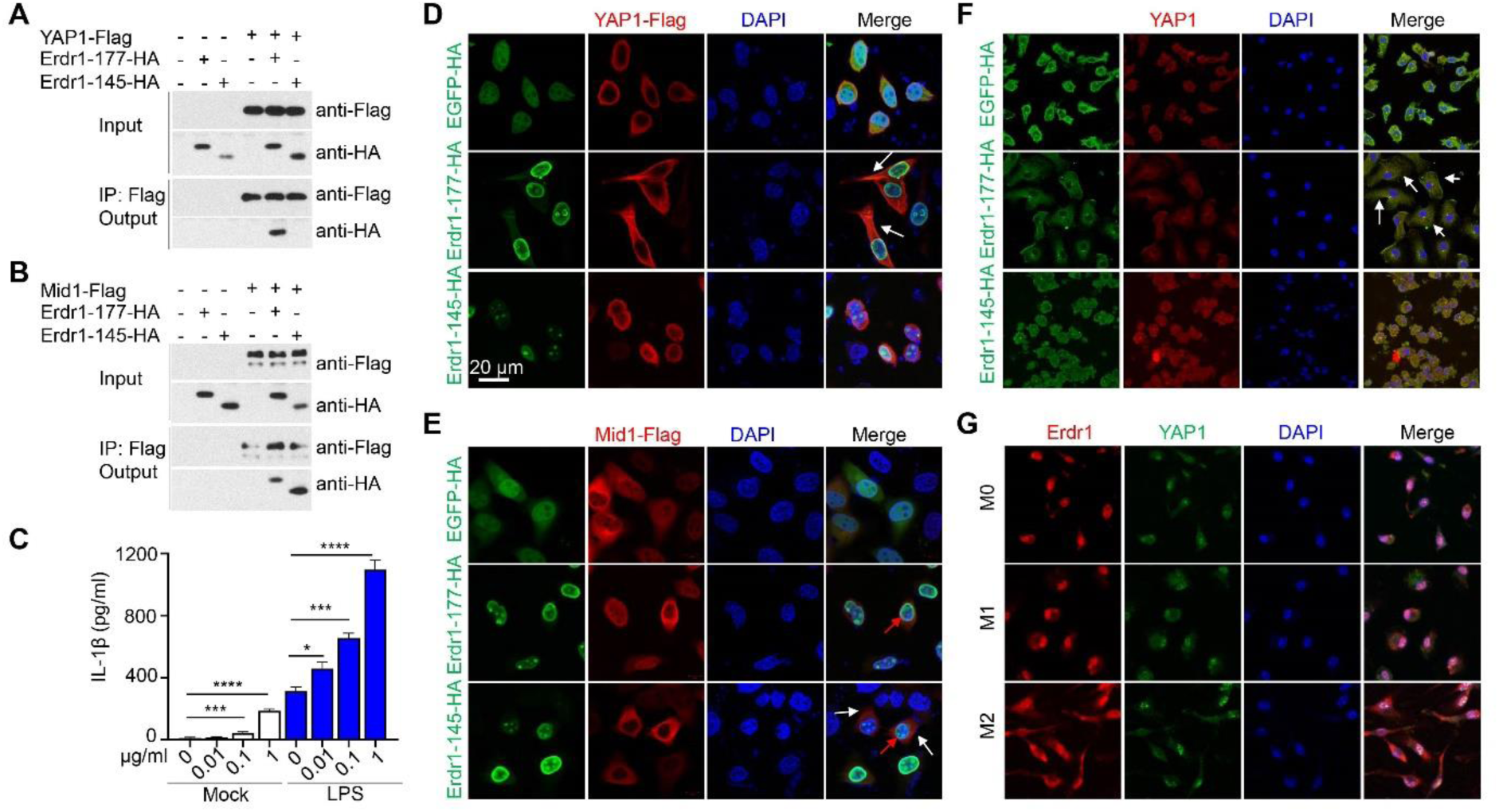
Erdr1 displays distinct interaction with YAP1 and Mid1 by different domains and mediates macrophage polarization. **(A-B)** Co-immunoprecipitation (Co-IP) and immunoblot analysis. Flag tag protein: YAP-Flag (**A**), Mid1-Flag (**B**), and HA-tag protein (Erdr1-177-HA and Erdr1-145-HA) were co-expressed in 293 T cells for 24 hours and cell lysates were subjected to IP using Flag beads. Both inputs and Co-IP fractions (Output) were subjected to Immunoblotting with anti-Flag or anti-HA antibodies as indicated. Erdr1-177, Full-length Erdr1; Erdr1-145, Deficiency in c-terminal 32 amino acid domain. Flag tag and HA tag were added at the C-terminal of the gene. **(C)** Detecting IL-1β production at supernatant with the indicated concentration of Erdr1-145 treatment in BMDM without or with 1 μg/ml LPS stimulation for 24 hours (n=4). **(D)** Co-overexpression of YAP1-Flag with Erdr1-177-HA or Erdr1-145-HA in Hela cell and fluorescence staining using anti-HA and anti-Flag antibodies. White arrows point to the direct interaction of Erdr1-177-HA with YAP1-Flag in the cytoplasm. **(E)** Co-overexpression of Mid1-Flag with Erdr1-177-HA or Erdr1-145-HA in Hela cell and fluorescence staining using anti-HA and anti-Flag antibodies. Red arrows point to the direct interaction of Mid1-Flag and Erdr1-177-HA/Erdr1-145-HA at the nuclear membrane. White arrows indicated Erdr1-145 promoted Mid1 localized at cytoplasm. **(F)** Immunofluorescence staining of overexpressed Erdr1-177-HA or Erdr1-145-HA by anti-HA antibody and endogenous YAP1 in BMDM. White arrows indicated the direct interaction of Erdr1-177 with YAP1 in the cytoplasm. **(G)** Immunofluorescence staining of endogenous Erdr1 and YAP1 in M0, M1, and M2 macrophages. Scale bar was marked in (**D**), it in represents 20 μm. Data represent mean ± SD. *p<0.05, ***p<0.001, ****p<0.001. Data are representative of two independent experiments.

**Figure 6.**
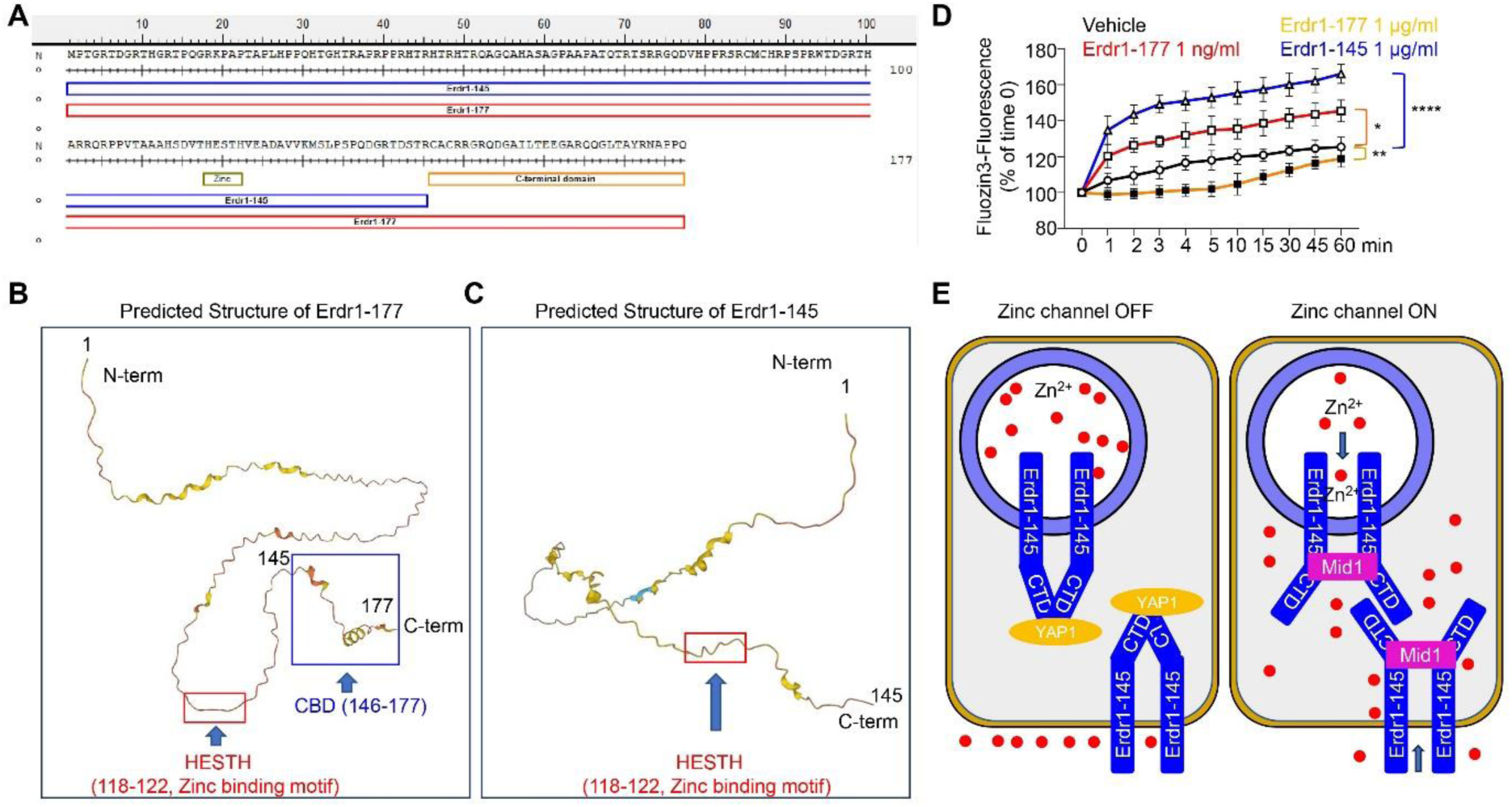
Erdr1 is a potent zinc channel by displaying different conformation. **(A)** Full-length Erdr1 (Erdr1-177) and C-terminal 32 amino acid deficiency Erd1 (Erdr1-145) amino acid sequence. There is a potent zinc binding motif HESTH (118-122). The C-terminal domain indicates the YAP1 binding domain. **(B-C)** AlphaFold structure prediction of Erdr1-177 (https://alphafold.ebi.ac.uk/entry/O88727) (**B**) and Erdr1-145 (https://alphafold.ebi.ac.uk/entry/Q6PED5) (**C**). The potent zinc-binding motif HEXXH (118-122) displayed hidden in Erdr1-177 but was exposed in Erdr1-145. **(D)** Detecting the intracellular zinc mobilization with zinc indicator. RAW-246.7 cell which was cultured in 96 wells plate with 0.6*10^6^ cell/ml were treated with 1 μg/ml LPS, 10 μM ZnCl2, indicated Erdr1 protein, and 1 μM FluoZin-3-AM in serum-free medium. The displayed values represent fluorescence (Ex/Em, 494/516) normalized to the initial time point (time 0). The p values were determined by two-way ANOVA. *p<0.05, **p<0.01, ****p<0.001. One representative experiment of mean ± SD of n = 4 experiments is shown. **(E)** Working model of Erdr1 acts as a zinc channel. Zinc channel off achieved by Erdr1-YAP1 display. Zinc channel on achieved by Erdr1-Mid1 interplay.

Using AlphaFold protein structure prediction, we revealed that Erdr1-177 and Erdr1-145 exhibit distinct conformations (Fig. 6B and 6C). Erdr1 harbors the HESTH sequence (118-122) (Fig. 6A, 6B and 6C), it belongs to a zinc-binding motif HEXXH, which is a common feature of many zinc metalloproteinases (57) and some members of the ZIP family of zinc transporters (66). The zinc binding site was hidden in the loop structure of Erdr-177 (Fig. 6B), but it exposed when C-terminal 32 amino acid deficiency (Erdr-145) (Fig. 6C). By using intracellular zinc indicator, we observed increased intracellular free zinc induced by LPS (Fig. 6D), which is important for pro-inflammatory cytokines production (67).The zinc influx could enhance by Erdr1-145, or low concentration of Erdr1-177 (1 ng/ml), but could suppress by high concentration of Erdr1-177 (1 μg/ml) in macrophages (Fig. 6D). This supported that Erdr1-Mid1 might act as the zinc channel for priming down-stream signaling (Fig. 6E). According to our hypothesis, by concentration change and the concomitant alterations in conformation, Erdr1 has different affinity with YAP1 and Mid1. Erdr1-YAP1 high affinity or, Erdr1-Mid1 high affinity, drive and polarize macrophage to two extremely inflammation states. Erdr1-177 (high concentration) which interacts with YAP1 can be considered to harbor anti-inflammatory conformation. Erdr1-145, which interacts with Mid1, can be considered to display pro-inflammatory conformation. Erdr1-177 and Erdr1-145 potentially can drive macrophage polarize into M2 and M1 states, respectively. To further investigate this hypothesis, we employed purified Erdr1-177 and Erdr1-145 proteins to assess their roles in mediating macrophage polarization. Our findings revealed that high concentration of Erdr1-177 (1 μg/ml) could spontaneously induce the expression of M2 marker genes, such as IL-10 and TGFβ (Supplemental Fig. 2A). Conversely, high concentration of Erdr1-145 (1 μg/ml) induced the expression of IL-6 and TNF, which are marker genes for M1 macrophages (Supplemental Fig. 2B). Notably, we observed that Erdr1-145 exhibits a dose-dependent effect on priming IL-1β production and could enhance LPS-induced IL-1β production (Fig. 5C). Overall, these data provide support for the notion that Erdr1 has the potential to act as a suppressor or an amplifier in inflammation response through conformational changes.

To investigate the interactions between Erdr1 and YAP1, as well as Erdr1 and Mid1 at the cellular level, we conducted experiments involving the overexpression of Erdr1-177 or Erdr1-145 along with YAP1 or Mid1 in Hela cells, with HA-tags and Flag-tags added at the C-terminus as indicated (Fig. 5D and Fig. 5E). Notably, Erdr1-177-HA co-localized specifically with YAP1-Flag in the cytoplasm, suggesting a direct interaction between these two proteins. Consistent with our Co-IP data, we did not observe a direct interaction between Erdr1-145 and YAP1 (Fig. 5D). On the other hand, we observed that both Erdr1-177 and Erdr1-145 interact with Mid1 at the nuclear envelope. Notably, Erdr1-145, but not Erdr1-177, demonstrated the ability to activate Mid1 and induce its translocation to the cytoplasm (Fig. 5E). In contrast, Erdr1-177 promoted the nuclear localization of Mid1, which represents the inhibitory state of Mid1 (Fig. 5E). These data highlight the distinct roles of different Erdr1 conformations in regulating YAP1 and Mid1, with Erdr1-177 licensing YAP1 OFF/Mid OFF, while Erdr1-145 forcing Mid1 ON/YAP1 ON.

We further explored the impact of Erdr1-177 and Erdr1-145 on the subcellular localization of endogenous YAP1 in macrophages. BMDM were overexpressed with Erdr1-177 or Erdr1-145 by lentivirus transduction. We observed that the overexpression of Erdr1-177 results in the sequestration of YAP1 in the cytoplasm (Fig. 5F) and induces a significant upregulated of IL-10 and TGF-β mRNA expression (Supplemental Fig. 2C), suggesting Erdr1-177 overexpression mediates macrophage polarization towards the M2 phenotype. However, overexpression of Erdr1-145 results in the nuclear localization of YAP1 (Fig. 5F) and a significant upregulation in the expression of IL-6 and TNF (Supplemental Fig. 2D), indicating that overexpression of Erdr1-145 drives macrophage polarization towards the M1 phenotype. These findings provide substantial evidence that Erdr1-177 polarizes macrophages towards the M2 state by regulating YAP1 OFF, while Erdr1-145 polarizes macrophages towards the M1 state by regulating YAP1 ON.

Furthermore, we illustrated the dynamic co-localization of endogenous Erdr1 and YAP1 in macrophages. Through immunofluorescence analysis, we detected a robust interaction between Erdr1 and YAP1 in the cytoplasm of M2 macrophages, whereas such interaction was not observed in M1 macrophages (Fig. 5G). These finding exhibit a distinct interaction pattern between endogenous Erdr1 and YAP1 in M1 and M2 macrophages.

Collectively, these observations strongly support our hypothesis that the dynamic interaction of Erdr1 with YAP1 and Mid1 drives and coordinates macrophage polarization.

### Erdr1 orchestrates macrophage functional and metabolic reprogramming and determines cell fate

Macrophage polarization is a vital functional response and adaptation to microenvironmental stimuli (60, 61). Activated macrophage not only exhibits distinct functions but also involves metabolic reprogramming (68) and ultimately have different cell death patterns (69–71). To further elucidate the role of Erdr1 in macrophage polarization, we further performed comprehensive analysis of Erdr1 role in inflammation, metabolic and cell death phenotype.

Erdr1-177 and Erdr1-145 protein, which represent different conformation of Erdr1, were observed to play distinct roles in regulating IL-1β production at indicated doses (1 μg/ml) with optimal cell density (0.6*10^6^ cell/ml) (Fig. 7A and 7B). Erdr1-177 inhibits IL-1β production upon LPS stimulation at mRNA level (Fig. 7A) and protein level in supernatant (Fig. 7B), while Erdr1-145 amplifies basal and LPS-induced IL-1β production (Fig. 7A and Fig. 7B). We validated these findings using immunoblotting analysis, manifesting Erdr1-177 inhibits pro-IL-1β cleaved to matured IL-1β, whereas Erdr1-145 promotes IL-1β maturation (Fig. 7D). Furthermore, we found Erdr1-177 inhibits Mid1 and YAP1, but Erdr1-145 promotes Mid1 and YAP1 expression (Fig. 7C and 7D), manifesting that Mid1 and YAP1 are also distinctly regulated by different Erdr1 conformations. These findings provide strong evidence that the highly dynamic fluctuations in conformation of Erdr1 have the capacity to dynamically regulate the levels of IL-1β, YAP1, and Mid1.

**Figure 7.**
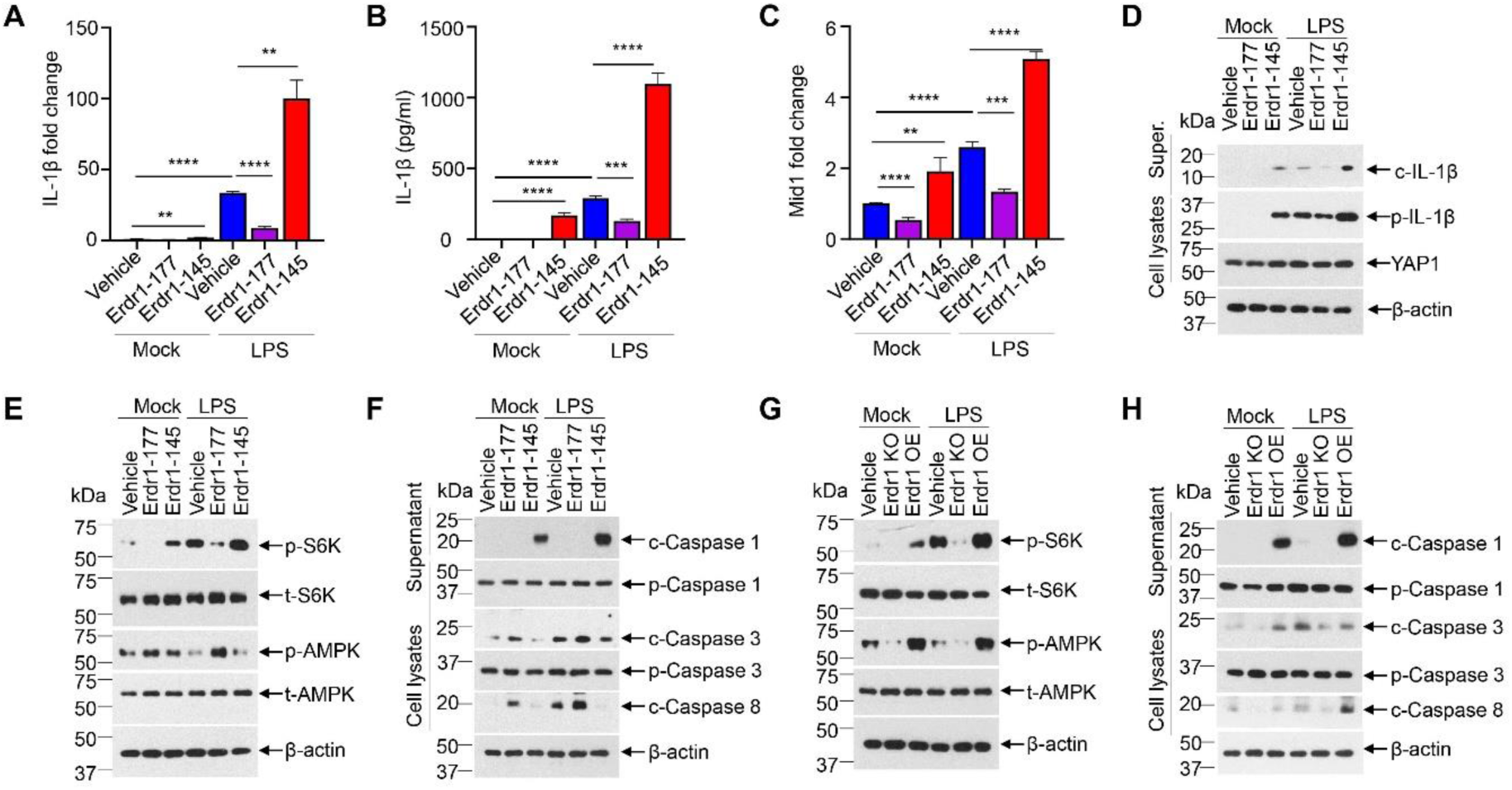
Erdr1 mediates macrophage functional and metabolic polarization and determines cell death phenotypes. **(A-F)** BMDM were treated as follows for 24 hours: 1. Vehicle; 2. Erdr1-177 (1μg/ml) 3. Erdr1-145 (1μg/ml); 4. Vehicle+ LPS (1μg/ml); 5. Erdr1-177 (1μg/ml) + LPS (1μg/ml); 6. Erdr1-145 (1μg/ml) + LPS (1μg/ml), Cell lysates and supernatant were collected for the following analysis. **(A)** qPCR analysis of IL-1β expression in the cell lysates (n=4). **(B)** Detecting IL-1β production at supernatant by ELISA (n=4). **c,** qPCR analysis of Mid1 expression in the cell lysates. (**A**) and (**C**) qPCR normalized to β-actin expression. **(D)** Immunoblot analysis of IL-1β, YAP1, and Mid1 levels in cell lysates and IL-1β in the supernatant; p-IL-1β, pro-IL-1β; c-IL-1β, cleaved-IL-1β. **(E)** Immunoblot analysis of key proteins involved in metabolism as indicated. (**F)** Immunoblot analysis of key proteins involved in apoptosis and pyroptosis as indicated. (**G-H)** BMDM were genetic knockout of Erdr1 (Erdr1 KO) or overexpressed Erdr1 (Erdr1 OE) by lentivirus transduction, the vehicle represents transduction with vehicle control lentivirus. Cell lysates were collected for immunoblot analysis with indicated antibodies without and with 1μg/ml LPS treatment for 24 hours. p-S6K, phospho-S6K; t-S6K, total S6K; p-AMPK, phospho-AMPK; t-AMPK, total AMPK; p-Caspase1, pro-caspase1; c-Caspase1, cleaved caspase1; p-Caspase3, pro-caspase3; c-caspase3, cleaved caspase3; c-caspase8, cleaved caspase8. (**A-H)** Cell density is 0.6*10^6^/ml and seeded in 24 well plates. Data represent mean ± SD. *p<0.05, **p<0.01, ***p<0.001 and, ****p<0.0001. Data are representative of three independent experiments

We then investigated if Erdr1 plays a role in mediating metabolic transition. Macrophage M1 polarization is characterized by the activation of mTORC1-S6K signaling (68), which represents an anabolic state. In contrast, macrophage M2 polarization is characterized by the activation of AMPK signaling (68), representing a catabolic state. Our findings demonstrated that Erdr1-177 protein inhibits p-S6K activation while promotes p-AMPK activation in basal and LPS-stimulated macrophages (Fig. 7E). Conversely, Erdr1-145 protein strikingly promotes p-S6K activation but inhibits p-AMPK activation in basal and LPS stimulated macrophage, compared to vehicle control (Fig. 7E). These data support that Erdr1 has the potential to polarized macrophage metabolism to pro-anabolic state or pro-catabolic state by conformation changes. Furthermore, we observed that Erdr1 deficiency macrophage exhibits low basal and LPS-induced p-S6K and p-AMPK activation, whereas Erdr1 overexpression macrophage displays high basal and LPS- induced p-S6K and p-AMPK activation (Fig. 7G), suggesting Erdr1 is essential both for anabolic and catabolic metabolism. These observations indicate that Erdr1 decides the metabolic intensity, phenotype, and transition within macrophages via concentration and conformational changes.

Furthermore, distinct activated macrophages undergo different cell death phenotype. M2 macrophage undergo apoptotic death, which elicit anti-inflammatory response. By contrast, M1 macrophage undergo pyroptosis, which is a proinflammatory form of cell death (69–71). Our data show that both Erdr1-177 and Erdr1-145 protein supplementation promote unstimulated macrophages death (Supplemental Fig. 2E). However, Erdr1-177 protects LPS-induced macrophage from death but Erdr1-145 aggravates LPS-induced cell death (Supplemental Fig. 2E). Our data further demonstrated that Erdr1-177 promotes apoptosis by enhancing the activation of caspase3 and caspase8, while inhibiting pyroptosis by reducing caspase1 activation in macrophages (Fig. 7F), indicating Erdr1-177 acts as a pro-apoptotic factor. Conversely, Erdr1-145 inhibits apoptosis but promotes pyroptosis in unstimulated and LPS-stimulated macrophages (Fig. 7F), supporting Erdr1 has the potential to be a pro-pyroptotic factor. Furthermore, we observed Erdr1 knockout reduces, whereas Erdr1 overexpression enhances both caspase3/8 and caspase1 activation in macrophage compared to control (Fig. 7H), implying high level of Erdr1 is essential for caspase-dependent cell death process. Even lack of caspase-dependent programmed cell death, we still found that Erdr1 deficiency in macrophages result in reduced cell viability (Supplemental Fig. 2F). It is not hard to understand this considering both catabolic and anabolic metabolism are blocked in Erdr1 deficiency macrophage (Fig. 7G), indicating basal expression of Erdr1 is essential for cell survive. In summary, these findings suggest that a narrow range of Erdr1 mediates cell survive, otherwise dysregulated Erdr1 determines macrophage programmed cell death through either a pro-apoptotic or pro-pyroptotic pathway.

Taken together, these data reveal dynamic change of Erdr1 also mediates metabolic transitions and determines cell fate.

## Discussion

This study aims to elucidate the function and mechanism of Erdr1 in IL-1β production and macrophage polarization. Erdr1 was discovered to exhibit a negative correlation with IL-18 (62) and purified Erdr1 protein played the anti-inflammatory role by inhibiting the production of various cytokines such as TNF, IL-6, and IL-1β (8, 63). Our data also manifest the anti-inflammatory role of Erdr1. What interesting is we uncover that Erdr1 has the potential to display pro-inflammatory role which highly associated with Erdr1 dose and cell densities. At optimal cell density, both endogenous and exogenous Erdr1 plays role in IL-1β production following a bell-shaped model. This Erdr1 dose-dependent threshold model has also been noted in Erdr1-mediated metastasis in murine embryonic fibroblasts (MEF) (35, 72), indicating Erdr1 has highly dynamic functions.

We evidence that the dual role of Erdr1 in IL-1β production is achieved by intricate Erdr1 level which displayed differently in M1/M2 macrophages. We found Erdr1 was significantly upregulated in M2 macrophages, and might act as a M2 marker gene, which is consistent with previous research (73). It is worth noting that we observed Erdr1 was dramatically suppressed in M1 macrophages. Down-regulated Erdr1 was also observed in other immune and non-immune cells upon activation. Erdr1 exhibited downregulation in CD4 T cells upon TCR stimulation (5). Erdr1 was also downregulated in tissues of patients with inflammation disorder disease (9, 10, 63), tumor cell (62), and was remarkably inhibited in bleomycin-induced fibroblast (56). These support that reduced Erdr1 acts to activate specific functions of the cell. Our data manifest that decreased Erdr1 undergoes conformation change and promotes IL-1β production and drives the macrophage pro-inflammatory polarization, whereas elevated Erdr1 inhibits IL-1β production and drives the macrophage anti-inflammatory polarization.

We figure out Erdr1 drives macrophage polarization by distinct interaction with YAP1 and Mid1. Consistent with previous research (24, 26, 55), we observed the significantly up-regulated expression of YAP1 and Mid1 in pro-inflammatory macrophages. Notably, YAP1 and Mid1 are both essential for Erdr1-mediated LPS-induced IL-1β production. We reveal Erdr1 acts as an upstream regulator of YAP1 and Mid1, regulates their expression, subcellular localization, and downstream gene expression, which includes IL-1β. Additionally, YAP1 and Mid1 have also been identified as promoters of fibrosis (55, 64). Down-regulated Erdr1 was also reported to involve in the pathogenesis of bleomycin-induced fibrosis in fibroblasts (56). CTGF, the central mediator of fibrosis (74) and a signature gene of YAP1 and Mid1 signaling (55, 65), is significantly regulated by Erdr1 (Fig. 4J). This indicates by regulating YAP1 and Mid1, Erdr1 also regulates fibrosis pathogenesis. Except for the role of Erdr1 in IL-1β, we also reveal Erdr1 distinctly drive M2 maker gene (IL-10 and TNFβ) or other M1 marker gene (IL-6 and TNF) expression. Our data demonstrate that Erdr1 acts as the driving force for macrophage functional polarization: 1. Erdr1-YAP1 interaction drives anti-inflammatory M2 state; 2. Erdr1-Mid1 interaction drives pro-inflammatory M1 state (Fig. 8A).

**Fig. 8.**
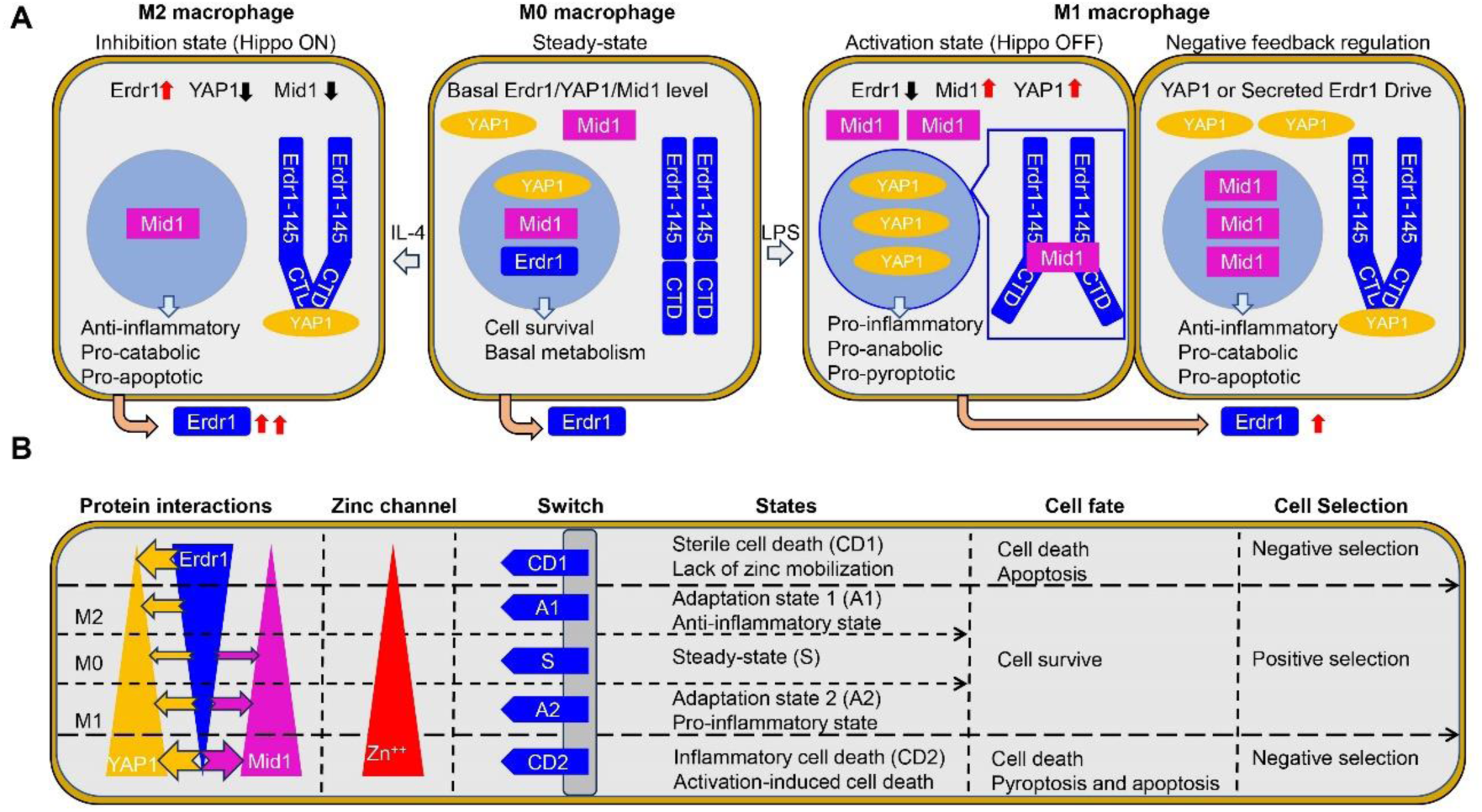
Working model of Erdr1 in macrophages polarization and cell fate determination. **(A)** Erdr1 drives macrophage polarization. Erdr1 has concentration and conformation change during macrophage polarization and plays multifunction in M1 and M2 macrophage. 1. The M0 macrophage represents the steady state of macrophage with basal levels of Erdr1, Mid1, and YAP1 expression, accompanied by a basal metabolism level. Erdr1 acts as a cell survival factor at this state. 2. M0 macrophages are induced to M2 macrophage with 20 ng/ml IL-4 treatment for 24 hours. In M2 macrophages, Erdr1 is endogenously upregulated, with increased cytoplasm localization and increased secretion to the extracellular matrix. The upregulated Erdr1 interacts with YAP1 through its C-terminal domain (Erdr1-CBD), which consists of 32 amino acids. This interaction sequesters YAP1 in the cytoplasm and reduces YAP1 expression (Hippo ON), consequently leading to Mid1 inactivation. Erdr1 acts as an anti-inflammatory, pro-catabolic and pro-apoptotic factor in M2 state. 3. M1 macrophages are induced from M0 macrophages by 1 μg/ml LPS treatment for 24 hours. In M1 macrophages, Erdr1 is endogenously downregulated and increased nuclear localization but increased release to extracellular matrix. The downregulated Erdr1 undergoes conformation shift and disconnecting with YAP1 but exhibits a high affinity with Mid1 through its Erdr1-145 domain (Erdr1-145, or Erdr1Δ32) at the nuclear envelope and forces YAP1 ON. Erdr1 acts as a pro-inflammatory, pro-anabolic and pro-pyroptotic factor. Additionally, negative feedback regulation can be initiated in M1 macrophage by up-regulated endogenous YAP1 and increased accumulation of secreted Erdr1. This figure illustrates the main expression level (red arrow indicates upregulated, black arrows indicates downregulated), subcellular localization, and dynamic interaction of Erdr1 with Mid1and YAP1 in M0, M1 and M2 macrophages. **(B)** working model of Erdr1 in cell fate determination. The interplay of Erdr1 with Mid1 and YAP1 shape macrophage heterogeneity and determines the cell survival or death, and the death patterns.

Metabolic reprogramming stands as a pivotal hallmark of macrophage polarization. YAP1 and Mid1 have both been documented to enhance pro-anabolic response (18, 58, 75–77). Notably, Erdr1 has been implicated in YAP1 and Mid1-mediated metabolism (18, 38). YAP1 has been reported to be essential for mTOR signaling, and its deficiency in dental epithelial stem cells resulted in decreased Erdr1 expression, ultimately leading to the inhibition of mTOR signaling (18). Moreover, elevated levels of Erdr1 and Mid1 have been observed to participate in unique metabolism (dampened oxidative phosphorylation) in β-cells harboring the SLC30A8 (Zinc transporter) R138X mutation (38). These suggest Erdr1 might enhance mTOR signaling by working with YAP1 and Mid1, which is consistent with our observation that Erdr1 by activating YAP1 and Mid1 signaling, has the pro-anabolic capability by activating p-S6K. However, our data also manifest Erdr1 has the pro-catabolic potential by activating p-AMPK, when at a high concentration. This phenomenon was also observed in previous research in T cells when Erdr1 was upregulated to a high level (78). In fact, Erdr1 highly expressed T cells was found to have increased both glycolysis and oxidative phosphorylation (78), which are downstream of mTORC1 and p-AMPK respectively (68, 79), indicating Erdr1 might involve in both anabolic and catabolic metabolism. Our data for the first time demonstrate that Erdr1 is essential for both anabolic and catabolic metabolism by Erdr1 knockout and overexpression experiment (Fig.7G). However, the two different conformations of Erdr1 can only mediate the metabolic polarization to pro-catabolic or pro-anabolic state, with antagonize each other (Fig.7E). This represents by binding with YAP1 and Mid1 with different affinities, Erdr1 mediates metabolic polarized to different extreme states. Supporting this, β cell which has increased expression of Erdr1 and Mid1 exhibited increase insulin secretion but dampened oxidative phosphorylation (38). This further support Erdr1-Mid1 affinity has the potential to prime mTORC1-glycolysis but inhibit p-AMPK-oxidative phosphorylation pathway. Collectively, our data suggest Erdr1 mediates macrophage metabolic polarization: 1. Erdr1-YAP1 affinity primes pro-catabolic response; 2. Erdr1-Mid1 affinity drives pro-anabolic response (Fig. 8A). As we know, p-AMPK activation promotes macrophage M2 polarization (80), while mTOR-p-S6K activation promotes IL-1β production in M1 macrophages (81). Thus, these further reinforce the idea that Erdr1 mediates the macrophage polarization also by driving metabolic reprograming.

YAP1 was reported to play important roles in regulating cell death phenotypes (82, 83). Activated YAP1 restrains apoptosis (82, 83), but enhances inflammatory activation and promotes pyroptosis in macrophage (25). Conversely, YAP1 ablation intensifies caspase3/8/9-dependent cell death (84). Previous studies reported Erdr1 acts as a pro-apoptotic factor via inducing caspase3 activation (5, 85). Our data consistent with that and manifest Erdr1 by driving YAP1 OFF induces caspase3 and caspase8 activation and promotes apoptosis (Fig. 7F and 7H). Erdr1 has also been associated with caspase1 and involved in progressive degenerative disease (86). Our data for the first time figure out the pro-pyroptotic role of Erdr1 (Fig. 7F and 7H) by driving YAP1 ON. In Erdr1-deficient macrophages, both apoptosis and pyroptosis were inhibited (Fig. 7H), thereby illustrating the involvement of Erdr1 in both pro-apoptotic and pro-pyroptotic pathways. Erdr1 was observed to only mediate cell survival at a narrow range, otherwise cause rapid cell death (72) . We also observed this phenomenon. Erdr1 deficiency and overexpressed macrophages both elicit cell death (Supplemental Fig. 2F) but manifest distinct cell death patterns, with Erdr1 deficiency induces caspase-independent cell death, while Erdr1 overexpression causes caspase-dependent cell death (Fig. 7H). These support that Erdr1 is essential for programed cell death and determines cell fate: 1. Erdr1-YAP1 affinity induces pro-apoptotic response. 2. Erdr1-Mid1 affinity primes pro-pyroptotic activation. Additionally, negative feedback mechanisms exist by heightened YAP1 or elevated extracellular accumulation of Erdr1, which foster Erdr1-YAP1 interaction and consequently facilitate cells apoptotic death, act to alleviate the inflammation (Fig. 8A). We emphasize that the fine and dynamic interaction between Erdr1 and YAP1 and Mid1 play pivotal roles in facilitating cellular adaptation and survive. Otherwise, a polarized interaction with YAP1 or Mid1 triggers cellular demise and decides the pattern of cell death (Fig. 8B).

YAP1 serves as a downstream effector of the Hippo pathway which is a highly conserved signaling pathway that governs organ size through the regulation of cell proliferation, apoptosis, and stem cell self-renewal (27). Over the years, extensive research has revealed that the Hippo ON (YAP1 OFF) can be achieved through a sequential cascade of kinase activations (27), which including the formation of a complex between Mst1/2 kinases and SAV1 and subsequently phosphorylates and activates LATS1/2 kinases (27). However, different from LATS1/2 kinases dependent way in regulating YAP1, a unique YAP1 regulation were observed in Tregs cells which exhibited altered levels of Erdr1 and Mid1(19). Mechanical cues, including cell density which as the cell intrinsic mechanical cues, have consistently been shown to regulate Hippo-YAP signaling by mediating actin rearrangement (17, 25, 87–89). Erdr1 has been suggested to be involved in actin modulation (6) and cell density dependent concentration change (Fig. 3). It is highly likely that Erdr1 serves as the sensor and works with Mid1 for transducing mechanical cues into intracellular signaling and driving actin remodeling. This, in turn, regulates Hippo-YAP1 signaling. Our findings suggest that any factors capable of influencing the intricate concentration changes of Erdr1 can modulate Hippo-YAP1 signaling. Collectively, we have successfully demonstrated that Erdr1 acts as an upstream regulator of YAP1 through direct interaction. Most importantly, we found that the interaction with YAP1 is highly linked to Erdr1 concentration and concomitant conformational shifts. These findings suggest the critical role of Erdr1 in regulating YAP1 through the non-classical Hippo pathway.

However, the research progress of Erdr1 was hindered by several obstacles: Firstly, Erdr1 global knockout resulted in 100% embryonic lethality (5, 90), which limited research in mice. This also indicates Erdr1 plays a crucial role in cell physiological processes. Secondly, although Erdr1 has been mapped to sex chromosomes in mice, due to it has a high GC content (72) and contains many repeats (32), making identifying the entire genome sequence of Erdr1 unavailable until recently (33). Thirdly, although human Erdr1 mRNA and protein have been detected with 100% identical to mice Erdr1 (2, 72), as of now there is still lack of accessible human Erdr1 mRNA and protein information within the NCBI database, attributed to the incomplete sequencing of the Erdr1 localization region in humans (91), this hampers Erdr1 being identified as a pivotal target in humans for further research. There is an urgent need to map the localization of Erdr1 in the human genome and complete the gene sequencing. At the same time, it is believed that Erdr1 should be also exist and highly conserved in other species. Efforts to further explore Erdr1 across different species will contribute to better understanding the evolutionary process of life.

In this study, at the cellular level, we demonstrate that Erdr1 plays a crucial role in orchestrating macrophage functional and metabolic reprogramming and ultimately determining cell fate through dynamic interplay with YAP1 and Mid1 (Fig. 8). The function of Erdr1 depends on extracellular stimulus pattern, strength, and duration. Our research highlights the dynamic regulation of Erdr1 as the dominant factor for the macrophage spectrum oscillating between the M1 and M2 extreme polarized states (Fig. 8B). The working model of Erdr1 in macrophages suggests that Erdr1 is involved in both the positive and negative selection of the macrophage repertoire (Fig. 8B). Erdr1 as a potent zinc channel and mediates the intracellular zinc mobilization, which are crucial for macrophage pro-inflammatory activation (67, 92). Just set up the macrophage harbor an extremely high level of Erdr1, if there are no stimuli to decrease the Erdr1 level, the zinc channel is sustained turn off by Erdr1-YAP1 interplay, it means macrophage cannot prime pro-inflammatory response and undergoes apoptosis. If Erdr1 was dramatic decreased, the zinc channel is fully turn on by Erdr-Mid1 interplay, it represents macrophages was overactivated and resulted in activation-induced cell death. These events caused reduced life span of macrophage and dysregulated macrophage activation. If erdr1 level were manipulated to a sustained range which help macrophage escaped from these two types of cell death and undergo sustained proliferation or/and activation, it may cause myeloid leukemia. Only If Erdr1 level was proper induced, it mediates macrophage reprograming, shapes macrophage heterogeneity and help macrophage survival (Fig. 8B). Understanding the molecular mechanism of macrophage polarization orchestrated by Erdr1 stands to profoundly enhance our ability to therapeutically modulate inflammation more precisely and efficiently. Moreover, it’s crucial to highlight our significant observation that Erdr1 regulates YAP1 signaling through the non-classical Hippo pathway. These findings imply the critical role of Erdr1 in maintaining cellular and tissue homeostasis, extending beyond its impact on macrophages alone. Erdr1 (5), YAP1 (93) and Mid1 (76) have been demonstrated also distinctly induced during T cell polarization. The negative correlation of Erdr1 with YAP1/Mid1 in activated T cells also suggested the working model of Erdr1 with YAP1 and Mid1, at least, applicable to the T cell selection process.

Furthermore, Erdr1’s high degree of conservation across species and its importance in cellular metabolism and cell fate determination underscore the potential significance of this finding. Theoretically, Erdr1 acts as an inhibitor for cellular anabolic reactions by interplay with YAP1, while acts as an activator by interplay with Mid1. The working model of Erdr1 with YAP1 and Mid1 supports the cellular “reaction-diffusion” theory model proposed by Alan Mathison Turing in 1952 (94). This signifies that we have discovered the fundamental law of biology, marking a no doubt landmark progress in the field of theory biology. Practically, Erdr1 acts as the cell fate determiner, the dysregulated Erdr1 within cells is the root cause of metabolic shifts and cell fate alters, which are the hallmarks of the most common diseases (95). By modulating Erdr1, we can easily reverse unfavorable cell fates into favorable ones. Most importantly, extricate Erdr1 protein can manipulate the intricate Erdr1 level and finally determines cell fates. This represents we have mastered a simple approach to beat the most prevalent diseases, such as cancer, diabetes, auto-inflammatory diseases, and degenerative diseases.

## Authors’ Contributions

Y.W. (during pregnancy) developed the conception, designed, and performed the experiments. Y.W. interpreted the data and wrote the manuscript. Throughout the manuscript, the term “we” refers to Y.W. and her son, who actively participated in this research during his fetal development.

## Acknowledgments

This work is supported by the National Institutes of Health (NIH) Grant P01 AI060699. Sincerely thank Dr. Stanley Perlman from The University of Iowa for his significant contribution to the manuscript revision. Sincere appreciation to Dr. Ning Li from Beijing Capital Agribusiness Co., Ltd., for actively participating in the discussion and for contributing to the manuscript revision.

## Reference

1. Dormer, P., E. Spitzer, and W. Moller. 2004. EDR is a stress-related survival factor from stroma and other tissues acting on early haematopoietic progenitors (E-Mix). Cytokine 27: 47–57.

2. Dormer, P., E. Spitzer, M. Frankenberger, and E. Kremmer. 2004. Erythroid differentiation regulator (EDR), a novel, highly conserved factor I. Induction of haemoglobin synthesis in erythroleukaemic cells. Cytokine 26: 231–242.

3. Kim, M. S., S. Lee, S. J. Jung, S. Park, K. E. Kim, T. S. Kim, H. J. Park, and D. Cho. 2019. Erythroid differentiation regulator 1 strengthens TCR signaling in thymocytes by modulating calcium flux. Cell Immunol 336: 28–33.

4. Kim, M. S., D. Park, S. Lee, S. Park, K. E. Kim, T. S. Kim, H. J. Park, and D. Cho. 2022. Erythroid Differentiation Regulator 1 Strengthens TCR Signaling by Enhancing PLCgamma1 Signal Transduction Pathway. Int J Mol Sci 23.

5. Soto, R., C. Petersen, C. L. Novis, J. L. Kubinak, R. Bell, W. Z. Stephens, T. E. Lane, R. S. Fujinami, A. Bosque, R. M. O’Connell, and J. L. Round. 2017. Microbiota promotes systemic T-cell survival through suppression of an apoptotic factor. Proc Natl Acad Sci U S A 114: 5497–5502.

6. Lee, H. R., S. Y. Huh, D. Y. Hur, H. Jeong, T. S. Kim, S. Y. Kim, S. B. Park, Y. Yang, S. I. Bang, H. Park, and D. Cho. 2014. ERDR1 enhances human NK cell cytotoxicity through an actin-regulated degranulation-dependent pathway. Cell Immunol 292: 78–84.

7. Kim, M. S., S. Lee, S. Park, K. E. Kim, H. J. Park, and D. Cho. 2020. Erythroid Differentiation Regulator 1 Ameliorates Collagen-Induced Arthritis via Activation of Regulatory T Cells. Int J Mol Sci 21.

8. Kim, K. E., S. Kim, S. Park, Y. Houh, Y. Yang, S. B. Park, S. Kim, D. Kim, D. Y. Hur, S. Kim, H. J. Park, S. I. Bang, and D. Cho. 2016. Therapeutic effect of erythroid differentiation regulator 1 (Erdr1) on collagen-induced arthritis in DBA/1J mouse. Oncotarget 7: 76354–76361.

9. Kim, M., K. E. Kim, H. Y. Jung, H. Jo, S. W. Jeong, J. Lee, C. H. Kim, H. Kim, D. Cho, and H. J. Park. 2015. Recombinant erythroid differentiation regulator 1 inhibits both inflammation and angiogenesis in a mouse model of rosacea. Exp Dermatol 24: 680–685.

10. Kim, K. E., Y. Houh, J. Lee, S. Kim, D. Cho, and H. J. Park. 2016. Downregulation of erythroid differentiation regulator 1 (Erdr1) plays a critical role in psoriasis pathogenesis. Exp Dermatol 25: 570–572.

11. Zheng, D., T. Liwinski, and E. Elinav. 2020. Inflammasome activation and regulation: toward a better understanding of complex mechanisms. Cell Discov 6: 36.

12. Kelley, N., D. Jeltema, Y. Duan, and Y. He. 2019. The NLRP3 Inflammasome: An Overview of Mechanisms of Activation and Regulation. Int J Mol Sci 20.

13. Abo, H., B. Chassaing, A. Harusato, M. Quiros, J. C. Brazil, V. L. Ngo, E. Viennois, D. Merlin, A. T. Gewirtz, A. Nusrat, and T. L. Denning. 2020. Erythroid differentiation regulator-1 induced by microbiota in early life drives intestinal stem cell proliferation and regeneration. Nat Commun 11: 513.

14. Wu, J. M. F., Y. Y. Cheng, T. W. H. Tang, C. Shih, J. H. Chen, and P. C. H. Hsieh. 2018. Prostaglandin E(2) Receptor 2 Modulates Macrophage Activity for Cardiac Repair. J Am Heart Assoc 7: e009216.

15. Gumbiner, B. M., and N. G. Kim. 2014. The Hippo-YAP signaling pathway and contact inhibition of growth. J Cell Sci 127: 709–717.

16. Pavel, M., M. Renna, S. J. Park, F. M. Menzies, T. Ricketts, J. Fullgrabe, A. Ashkenazi, R. A. Frake, A. C. Lombarte, C. F. Bento, K. Franze, and D. C. Rubinsztein. 2018. Contact inhibition controls cell survival and proliferation via YAP/TAZ-autophagy axis. Nat Commun 9: 2961.

17. Zhao, B., X. Wei, W. Li, R. S. Udan, Q. Yang, J. Kim, J. Xie, T. Ikenoue, J. Yu, L. Li, P. Zheng, K. Ye, A. Chinnaiyan, G. Halder, Z. C. Lai, and K. L. Guan. 2007. Inactivation of YAP oncoprotein by the Hippo pathway is involved in cell contact inhibition and tissue growth control. Genes Dev 21: 2747–2761.

18. Hu, J. K., W. Du, S. J. Shelton, M. C. Oldham, C. M. DiPersio, and O. D. Klein. 2017. An FAK-YAP-mTOR Signaling Axis Regulates Stem Cell-Based Tissue Renewal in Mice. Cell Stem Cell 21: 91–106 e106.

19. Ni, X., J. Tao, J. Barbi, Q. Chen, B. V. Park, Z. Li, N. Zhang, A. Lebid, A. Ramaswamy, P. Wei, Y. Zheng, X. Zhang, X. Wu, P. Vignali, C. P. Yang, H. Li, D. Pardoll, L. Lu, D. Pan, and F. Pan. 2018. YAP Is Essential for Treg-Mediated Suppression of Antitumor Immunity. Cancer Discov 8: 1026–1043.

20. Hamon, A., D. Garcia-Garcia, D. Ail, J. Bitard, A. Chesneau, D. Dalkara, M. Locker, J. E. Roger, and M. Perron. 2019. Linking YAP to Muller Glia Quiescence Exit in the Degenerative Retina. Cell Rep 27: 1712–1725 e1716.

21. Gong, S., G. Cao, F. Li, Z. Chen, X. Pan, H. Ma, Y. Zhang, B. Yu, and J. Kou. 2021. Endothelial Conditional Knockdown of NMMHC IIA (Nonmuscle Myosin Heavy Chain IIA) Attenuates Blood-Brain Barrier Damage During Ischemia-Reperfusion Injury. Stroke 52: 1053–1064.

22. Lee, C. M., and J. Hu. 2013. Cell density during differentiation can alter the phenotype of bone marrow-derived macrophages. Cell Biosci 3: 30.

23. Vaughan-Jackson, A., S. Stodolak, K. H. Ebrahimi, E. Johnson, P. K. Reardon, M. Dupont, S. Zhang, J. S. O. McCullagh, and W. S. James. 2022. Density dependent regulation of inflammatory responses in macrophages. Front Immunol 13: 895488.

24. Zhou, X., W. Li, S. Wang, P. Zhang, Q. Wang, J. Xiao, C. Zhang, X. Zheng, X. Xu, S. Xue, L. Hui, H. Ji, B. Wei, and H. Wang. 2019. YAP Aggravates Inflammatory Bowel Disease by Regulating M1/M2 Macrophage Polarization and Gut Microbial Homeostasis. Cell Rep 27: 1176–1189 e1175.

25. Meli, V. S., H. Atcha, P. K. Veerasubramanian, R. R. Nagalla, T. U. Luu, E. Y. Chen, C. F. Guerrero-Juarez, K. Yamaga, W. Pandori, J. Y. Hsieh, T. L. Downing, D. A. Fruman, M. B. Lodoen, M. V. Plikus, W. Wang, and W. F. Liu. 2020. YAP-mediated mechanotransduction tunes the macrophage inflammatory response. Sci Adv 6.

26. Mia, M. M., D. M. Cibi, S. A. B. Abdul Ghani, W. Song, N. Tee, S. Ghosh, J. Mao, E. N. Olson, and M. K. Singh. 2020. YAP/TAZ deficiency reprograms macrophage phenotype and improves infarct healing and cardiac function after myocardial infarction. PLoS Biol 18: e3000941.

27. Ma, S., Z. Meng, R. Chen, and K. L. Guan. 2019. The Hippo Pathway: Biology and Pathophysiology. Annu Rev Biochem 88: 577–604.

28. Lu, T., R. Chen, T. C. Cox, R. X. Moldrich, N. Kurniawan, G. Tan, J. K. Perry, A. Ashworth, P. F. Bartlett, L. Xu, J. Zhang, B. Lu, M. Wu, Q. Shen, Y. Liu, L. J. Richards, and Z. Xiong. 2013. X-linked microtubule-associated protein, Mid1, regulates axon development. Proc Natl Acad Sci U S A 110: 19131–19136.

29. Collison, A. M., J. Li, A. P. de Siqueira, X. Lv, H. D. Toop, J. C. Morris, M. R. Starkey, P. M. Hansbro, J. Zhang, and J. Mattes. 2019. TRAIL signals through the ubiquitin ligase MID1 to promote pulmonary fibrosis. BMC Pulm Med 19: 31.

30. Chen, X., L. Wang, H. Yu, Q. Shen, Y. Hou, Y. X. Xia, L. Li, L. Chang, and W. H. Li. 2023. Irradiated lung cancer cell-derived exosomes modulate macrophage polarization by inhibiting MID1 via miR-4655-5p. Mol Immunol 155: 58–68.

31. Fang, M., A. Zhang, Y. Du, W. Lu, J. Wang, L. J. Minze, T. C. Cox, X. C. Li, J. Xing, and Z. Zhang. 2022. TRIM18 is a critical regulator of viral myocarditis and organ inflammation. J Biomed Sci 29: 55.

32. Chu, H. P., J. E. Froberg, B. Kesner, H. J. Oh, F. Ji, R. Sadreyev, S. F. Pinter, and J. T. Lee. 2017. PAR-TERRA directs homologous sex chromosome pairing. Nat Struct Mol Biol 24: 620–631.

33. Kasahara, T., K. Mekada, K. Abe, A. Ashworth, and T. Kato. 2022. Complete sequencing of the mouse pseudoautosomal region, the most rapidly evolving ‘chromosome’. bioRxiv: 2022.2003.2026.485930.

34. Armoskus, C., D. Moreira, K. Bollinger, O. Jimenez, S. Taniguchi, and H. W. Tsai. 2014. Identification of sexually dimorphic genes in the neonatal mouse cortex and hippocampus. Brain Res 1562: 23–38.

35. Mango, R. L., Q. P. Wu, M. West, E. C. McCook, J. S. Serody, and H. W. van Deventer. 2014. C-C chemokine receptor 5 on pulmonary mesenchymal cells promotes experimental metastasis via the induction of erythroid differentiation regulator 1. Mol Cancer Res 12: 274–282.

36. Kovačić, N. L. N. 2019. Mid1 is a novel mediator of subchondral bone resorption in antigen-induced arthritis. 4th annual meeting of the European Calcified Tissue Society, 11., Budapest, Hungary.

37. Perez, E. C., P. Xander, M. F. L. Laurindo, E. B. R. R. Novaes, B. C. Vivanco, R. A. Mortara, M. Mariano, J. D. Lopes, and A. C. Keller. 2017. The axis IL-10/claudin-10 is implicated in the modulation of aggressiveness of melanoma cells by B-1 lymphocytes. PLoS One 12: e0187333.

38. Kleiner, S., D. Gomez, B. Megra, E. Na, R. Bhavsar, K. Cavino, Y. Xin, J. Rojas, G. Dominguez-Gutierrez, B. Zambrowicz, G. Carrat, P. Chabosseau, M. Hu, A. J. Murphy, G. D. Yancopoulos, G. A. Rutter, and J. Gromada. 2018. Mice harboring the human SLC30A8 R138X loss-of-function mutation have increased insulin secretory capacity. Proc Natl Acad Sci U S A 115: E7642–E7649.

39. Verhagen, A. M., C. A. de Graaf, T. M. Baldwin, A. Goradia, J. E. Collinge, B. T. Kile, D. Metcalf, R. Starr, and D. J. Hilton. 2012. Reduced lymphocyte longevity and homeostatic proliferation in lamin B receptor-deficient mice results in profound and progressive lymphopenia. J Immunol 188: 122–134.

40. Sferruzzi-Perri, A. N., A. M. Macpherson, C. T. Roberts, and S. A. Robertson. 2009. Csf2 null mutation alters placental gene expression and trophoblast glycogen cell and giant cell abundance in mice. Biol Reprod 81: 207–221.

41. Laufer, B. I., K. Neier, A. E. Valenzuela, D. H. Yasui, R. J. Schmidt, P. J. Lein, and J. M. LaSalle. 2022. Placenta and fetal brain share a neurodevelopmental disorder DNA methylation profile in a mouse model of prenatal PCB exposure. Cell Rep 38: 110442.

42. Ratliff, W. A., D. Qubty, V. Delic, C. G. Pick, and B. A. Citron. 2020. Repetitive Mild Traumatic Brain Injury and Transcription Factor Modulation. J Neurotrauma 37: 1910–1917.

43. Jackson, I. L., F. Baye, C. P. Goswami, B. P. Katz, A. Zodda, R. Pavlovic, G. Gurung, D. Winans, and Z. Vujaskovic. 2017. Gene expression profiles among murine strains segregate with distinct differences in the progression of radiation-induced lung disease. Dis Model Mech 10: 425–437.

44. Descalzi, G., V. Mitsi, I. Purushothaman, S. Gaspari, K. Avrampou, Y. E. Loh, L. Shen, and V. Zachariou. 2017. Neuropathic pain promotes adaptive changes in gene expression in brain networks involved in stress and depression. Sci Signal 10.

45. Zhao, Z., T. Miki, A. Van Oort-Jansen, T. Matsumoto, D. S. Loose, and C. C. Lee. 2011. Hepatic gene expression profiling of 5’-AMP-induced hypometabolism in mice. Physiol Genomics 43: 325–345.

46. Hou, Y. J., R. Banerjee, B. Thomas, C. Nathan, A. Garcia-Sastre, A. Ding, and M. B. Uccellini. 2013. SARM is required for neuronal injury and cytokine production in response to central nervous system viral infection. J Immunol 191: 875–883.

47. Qiao, X., J. Y. Lu, and S. L. Hofmann. 2007. Gene expression profiling in a mouse model of infantile neuronal ceroid lipofuscinosis reveals upregulation of immediate early genes and mediators of the inflammatory response. BMC Neurosci 8: 95.

48. Vijay, R., A. R. Fehr, A. M. Janowski, J. Athmer, D. L. Wheeler, M. Grunewald, R. Sompallae, S. P. Kurup, D. K. Meyerholz, F. S. Sutterwala, S. Narumiya, and S. Perlman. 2017. Virus-induced inflammasome activation is suppressed by prostaglandin D(2)/DP1 signaling. Proc Natl Acad Sci U S A 114: E5444–E5453.

49. Li, W., M. J. Hofer, P. Songkhunawej, S. R. Jung, D. Hancock, G. Denyer, and I. L. Campbell. 2017. Type I interferon-regulated gene expression and signaling in murine mixed glial cells lacking signal transducers and activators of transcription 1 or 2 or interferon regulatory factor 9. J Biol Chem 292: 5845–5859.

50. Karlsen, T. R., M. B. Olsen, X. Y. Kong, K. Yang, A. Quiles-Jimenez, P. Kroustallaki, S. Holm, G. T. Lines, P. Aukrust, T. Skarpengland, M. Bjoras, T. B. Dahl, H. Nilsen, I. Gregersen, and B. Halvorsen. 2022. NEIL3-deficient bone marrow displays decreased hematopoietic capacity and reduced telomere length. Biochem Biophys Rep 29: 101211.

51. Rubin, C. M., D. A. van der List, J. M. Ballesteros, A. V. Goloshchapov, L. M. Chalupa, and B. Chapman. 2011. Mouse mutants for the nicotinic acetylcholine receptor ss2 subunit display changes in cell adhesion and neurodegeneration response genes. PLoS One 6: e18626.

52. Gangaplara, A., C. Martens, E. Dahlstrom, A. Metidji, A. S. Gokhale, D. D. Glass, M. Lopez-Ocasio, R. Baur, K. Kanakabandi, S. F. Porcella, and E. M. Shevach. 2018. Type I interferon signaling attenuates regulatory T cell function in viral infection and in the tumor microenvironment. PLoS Pathog 14: e1006985.

53. Pagliari, S., V. Vinarsky, F. Martino, A. R. Perestrelo, J. Oliver De La Cruz, G. Caluori, J. Vrbsky, P. Mozetic, A. Pompeiano, A. Zancla, S. G. Ranjani, P. Skladal, D. Kytyr, Z. Zdrahal, G. Grassi, M. Sampaolesi, A. Rainer, and G. Forte. 2021. YAP-TEAD1 control of cytoskeleton dynamics and intracellular tension guides human pluripotent stem cell mesoderm specification. Cell Death Differ 28: 1193–1207.

54. Mohseni, M., J. Sun, A. Lau, S. Curtis, J. Goldsmith, V. L. Fox, C. Wei, M. Frazier, O. Samson, K. K. Wong, C. Kim, and F. D. Camargo. 2014. A genetic screen identifies an LKB1-MARK signalling axis controlling the Hippo-YAP pathway. Nat Cell Biol 16: 108–117.

55. Chen, Q., C. Gao, M. Wang, X. Fei, and N. Zhao. 2021. TRIM18-Regulated STAT3 Signaling Pathway via PTP1B Promotes Renal Epithelial-Mesenchymal Transition, Inflammation, and Fibrosis in Diabetic Kidney Disease. Front Physiol 12: 709506.

56. Wang, Y., L. Zhang, T. Huang, G. R. Wu, Q. Zhou, F. X. Wang, L. M. Chen, F. Sun, Y. Lv, F. Xiong, S. Zhang, Q. Yu, P. Yang, W. Gu, Y. Xu, J. Zhao, H. Zhang, W. Xiong, and C. Y. Wang. 2022. The methyl-CpG-binding domain 2 facilitates pulmonary fibrosis by orchestrating fibroblast to myofibroblast differentiation. Eur Respir J 60.

57. Fukasawa, K. M., T. Hata, Y. Ono, and J. Hirose. 2011. Metal preferences of zinc-binding motif on metalloproteases. J Amino Acids 2011: 574816.

58. Posey, K. L., F. Coustry, A. C. Veerisetty, M. G. Hossain, M. J. Gambello, and J. T. Hecht. 2019. Novel mTORC1 Mechanism Suggests Therapeutic Targets for COMPopathies. Am J Pathol 189: 132–146.

59. Wright, P. E., and H. J. Dyson. 2015. Intrinsically disordered proteins in cellular signalling and regulation. Nat Rev Mol Cell Biol 16: 18–29.

60. Orecchioni, M., Y. Ghosheh, A. B. Pramod, and K. Ley. 2019. Macrophage Polarization: Different Gene Signatures in M1(LPS+) vs. Classically and M2(LPS-) vs. Alternatively Activated Macrophages. Front Immunol 10: 1084.

61. Murray, P. J. 2017. Macrophage Polarization. Annu Rev Physiol 79: 541–566.

62. Jung, M. K., Y. Park, S. B. Song, S. Y. Cheon, S. Park, Y. Houh, S. Ha, H. J. Kim, J. M. Park, T. S. Kim, W. J. Lee, B. J. Cho, S. I. Bang, H. Park, and D. Cho. 2011. Erythroid differentiation regulator 1, an interleukin 18-regulated gene, acts as a metastasis suppressor in melanoma. J Invest Dermatol 131: 2096–2104.

63. Houh, Y. K., K. E. Kim, H. J. Park, and D. Cho. 2016. Roles of Erythroid Differentiation Regulator 1 (Erdr1) on Inflammatory Skin Diseases. Int J Mol Sci 17.

64. Liu, F., D. Lagares, K. M. Choi, L. Stopfer, A. Marinkovic, V. Vrbanac, C. K. Probst, S. E. Hiemer, T. H. Sisson, J. C. Horowitz, I. O. Rosas, L. E. Fredenburgh, C. Feghali-Bostwick, X. Varelas, A. M. Tager, and D. J. Tschumperlin. 2015. Mechanosignaling through YAP and TAZ drives fibroblast activation and fibrosis. Am J Physiol Lung Cell Mol Physiol 308: L344–357.

65. Shome, D., T. von Woedtke, K. Riedel, and K. Masur. 2020. The HIPPO Transducer YAP and Its Targets CTGF and Cyr61 Drive a Paracrine Signalling in Cold Atmospheric Plasma-Mediated Wound Healing. Oxid Med Cell Longev 2020: 4910280.

66. Bin, B. H., J. Seo, and S. T. Kim. 2018. Function, Structure, and Transport Aspects of ZIP and ZnT Zinc Transporters in Immune Cells. J Immunol Res 2018: 9365747.

67. Haase, H., J. L. Ober-Blobaum, G. Engelhardt, S. Hebel, A. Heit, H. Heine, and L. Rink. 2008. Zinc signals are essential for lipopolysaccharide-induced signal transduction in monocytes. J Immunol 181: 6491–6502.

68. Kelly, B., and L. A. O’Neill. 2015. Metabolic reprogramming in macrophages and dendritic cells in innate immunity. Cell Res 25: 771–784.

69. Cong, Y., Y. Wang, T. Yuan, Z. Zhang, J. Ge, Q. Meng, Z. Li, and S. Sun. 2023. Macrophages in aseptic loosening: Characteristics, functions, and mechanisms. Front Immunol 14: 1122057.

70. Kloditz, K., and B. Fadeel. 2019. Three cell deaths and a funeral: macrophage clearance of cells undergoing distinct modes of cell death. Cell Death Discov 5: 65.

71. Robinson, N., R. Ganesan, C. Hegedus, K. Kovacs, T. A. Kufer, and L. Virag. 2019. Programmed necrotic cell death of macrophages: Focus on pyroptosis, necroptosis, and parthanatos. Redox Biol 26: 101239.

72. Mango, R. L. 2010. Stromal promotion of metastasis by erythroid differentiation regulator 1. University of North Carolina at Chapel Hill.

73. Lee, D. C., C. R. Ruiz, L. Lebson, M. L. Selenica, J. Rizer, J. B. Hunt, Jr., R. Rojiani, P. Reid, S. Kammath, K. Nash, C. A. Dickey, M. Gordon, and D. Morgan. 2013. Aging enhances classical activation but mitigates alternative activation in the central nervous system. Neurobiol Aging 34: 1610–1620.

74. Lipson, K. E., C. Wong, Y. Teng, and S. Spong. 2012. CTGF is a central mediator of tissue remodeling and fibrosis and its inhibition can reverse the process of fibrosis. Fibrogenesis Tissue Repair 5: S24.

75. Liu, E., C. A. Knutzen, S. Krauss, S. Schweiger, and G. G. Chiang. 2011. Control of mTORC1 signaling by the Opitz syndrome protein MID1. Proc Natl Acad Sci U S A 108: 8680–8685.

76. Boding, L., A. K. Hansen, G. Meroni, B. B. Johansen, T. H. Braunstein, C. M. Bonefeld, M. Kongsbak, B. A. Jensen, A. Woetmann, A. R. Thomsen, N. Odum, M. R. von Essen, and C. Geisler. 2014. Midline 1 directs lytic granule exocytosis and cytotoxicity of mouse killer T cells. Eur J Immunol 44: 3109–3118.

77. Matthes, F., M. M. Hettich, J. Schilling, D. Flores-Dominguez, N. Blank, T. Wiglenda, A. Buntru, H. Wolf, S. Weber, I. Vorberg, A. Dagane, G. Dittmar, E. Wanker, D. Ehninger, and S. Krauss. 2018. Inhibition of the MID1 protein complex: a novel approach targeting APP protein synthesis. Cell Death Discov 4: 4.

78. Muri, J., S. Heer, M. Matsushita, L. Pohlmeier, L. Tortola, T. Fuhrer, M. Conrad, N. Zamboni, J. Kisielow, and M. Kopf. 2018. The thioredoxin-1 system is essential for fueling DNA synthesis during T-cell metabolic reprogramming and proliferation. Nat Commun 9: 1851.

79. Wculek, S. K., G. Dunphy, I. Heras-Murillo, A. Mastrangelo, and D. Sancho. 2022. Metabolism of tissue macrophages in homeostasis and pathology. Cell Mol Immunol 19: 384–408.

80. Sag, D., D. Carling, R. D. Stout, and J. Suttles. 2008. Adenosine 5’-monophosphate-activated protein kinase promotes macrophage polarization to an anti-inflammatory functional phenotype. J Immunol 181: 8633–8641.

81. Kim, B., H. Y. Kim, B. R. Yoon, J. Yeo, J. In Jung, K. S. Yu, H. C. Kim, S. J. Yoo, J. K. Park, S. W. Kang, and W. W. Lee. 2022. Cytoplasmic zinc promotes IL-1beta production by monocytes and macrophages through mTORC1-induced glycolysis in rheumatoid arthritis. Sci Signal 15: eabi7400.

82. Cheng, Y., M. Mao, and Y. Lu. 2022. The biology of YAP in programmed cell death. Biomark Res 10: 34.

83. Zhang, X., A. Abdelrahman, B. Vollmar, and D. Zechner. 2018. The Ambivalent Function of YAP in Apoptosis and Cancer. Int J Mol Sci 19.

84. LeBlanc, L., B. K. Lee, A. C. Yu, M. Kim, A. V. Kambhampati, S. M. Dupont, D. Seruggia, B. U. Ryu, S. H. Orkin, and J. Kim. 2018. Yap1 safeguards mouse embryonic stem cells from excessive apoptosis during differentiation. Elife 7.

85. Kim, H. J., S. B. Song, Y. Yang, Y. S. Eun, B. K. Cho, H. J. Park, and D. H. Cho. 2011. Erythroid differentiation regulator 1 (Erdr1) is a proapototic factor in human keratinocytes. Exp Dermatol 20: 920–925.

86. Lee, J. H., C. H. Kim, D. G. Kim, and Y. S. Ahn. 2009. Microarray analysis of differentially expressed genes in the brains of tubby mice. Korean J Physiol Pharmacol 13: 91–97.

87. Petzold, J., and E. Gentleman. 2021. Intrinsic Mechanical Cues and Their Impact on Stem Cells and Embryogenesis. Front Cell Dev Biol 9: 761871.

88. Yu, F. X., B. Zhao, and K. L. Guan. 2015. Hippo Pathway in Organ Size Control, Tissue Homeostasis, and Cancer. Cell 163: 811–828.

89. Meng, K. P., F. S. Majedi, T. J. Thauland, and M. J. Butte. 2020. Mechanosensing through YAP controls T cell activation and metabolism. J Exp Med 217.

90. Zuo, E., Y. J. Cai, K. Li, Y. Wei, B. A. Wang, Y. Sun, Z. Liu, J. Liu, X. Hu, W. Wei, X. Huo, L. Shi, C. Tang, D. Liang, Y. Wang, Y. H. Nie, C. C. Zhang, X. Yao, X. Wang, C. Zhou, W. Ying, Q. Wang, R. C. Chen, Q. Shen, G. L. Xu, J. Li, Q. Sun, Z. Q. Xiong, and H. Yang. 2017. One-step generation of complete gene knockout mice and monkeys by CRISPR/Cas9-mediated gene editing with multiple sgRNAs. Cell Res 27: 933–945.

91. Rhie, A., S. Nurk, M. Cechova, S. J. Hoyt, D. J. Taylor, N. Altemose, P. W. Hook, S. Koren, M. Rautiainen, I. A. Alexandrov, J. Allen, M. Asri, A. V. Bzikadze, N. C. Chen, C. S. Chin, M. Diekhans, P. Flicek, G. Formenti, A. Fungtammasan, C. Garcia Giron, E. Garrison, A. Gershman, J. L. Gerton, P. G. S. Grady, A. Guarracino, L. Haggerty, R. Halabian, N. F. Hansen, R. Harris, G. A. Hartley, W. T. Harvey, M. Haukness, J. Heinz, T. Hourlier, R. M. Hubley, S. E. Hunt, S. Hwang, M. Jain, R. K. Kesharwani, A. P. Lewis, H. Li, G. A. Logsdon, J. K. Lucas, W. Makalowski, C. Markovic, F. J. Martin, A. M. Mc Cartney, R. C. McCoy, J. McDaniel, B. M. McNulty, P. Medvedev, A. Mikheenko, K. M. Munson, T. D. Murphy, H. E. Olsen, N. D. Olson, L. F. Paulin, D. Porubsky, T. Potapova, F. Ryabov, S. L. Salzberg, M. E. G. Sauria, F. J. Sedlazeck, K. Shafin, V. A. Shepelev, A. Shumate, J. M. Storer, L. Surapaneni, A. M. Taravella Oill, F. Thibaud-Nissen, W. Timp, M. Tomaszkiewicz, M. R. Vollger, B. P. Walenz, A. C. Watwood, M. H. Weissensteiner, A. M. Wenger, M. A. Wilson, S. Zarate, Y. Zhu, J. M. Zook, E. E. Eichler, R. J. O’Neill, M. C. Schatz, K. H. Miga, K. D. Makova, and A. M. Phillippy. 2023. The complete sequence of a human Y chromosome. Nature.

92. Rink, L., and H. Kirchner. 2000. Zinc-altered immune function and cytokine production. J Nutr 130: 1407S–1411S.

93. Stampouloglou, E., N. Cheng, A. Federico, E. Slaby, S. Monti, G. L. Szeto, and X. Varelas. 2020. Yap suppresses T-cell function and infiltration in the tumor microenvironment. PLoS Biol 18: e3000591.

94. Turing, A. M. 1990. The chemical basis of morphogenesis. 1953. Bull Math Biol 52: 153–197; discussion 119-152.

95. Schwartz, L., M. Henry, K. O. Alfarouk, S. J. Reshkin, and M. Radman. 2021. Metabolic Shifts as the Hallmark of Most Common Diseases: The Quest for the Underlying Unity. Int J Mol Sci 22.

