## Supplemental Figure 1 and 2 for "Erdr1 orchestrates macrophage polarization and determines cell fate via dynamic interplay with YAP1 and Mid1"

### Supplemental data

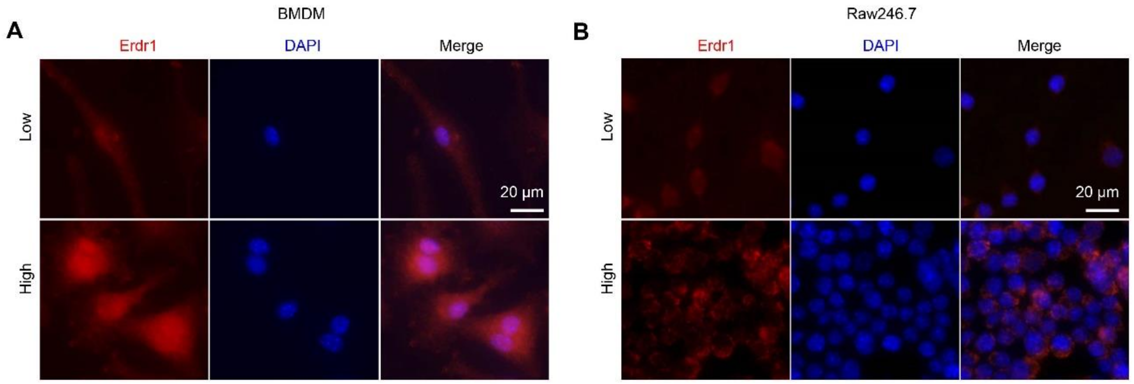


**Supplemental Figure 1 Cell density affects Erdr1 expression level and subcellular localization.**
BMDM (**A**) or RAW246.7 (**B**) Cell was seeded with low density (Low, 0.1*$10^6$ cells/ml) or high density (High, 1*$10^6$ cells/ml) in 24 well plates. Detecting Erdr1 subcellular localization at different cell densities by the Immunofluorescence staining of Erdr1 (Red) and counterstained by DAPI. Scale bar in represents 20 μm.

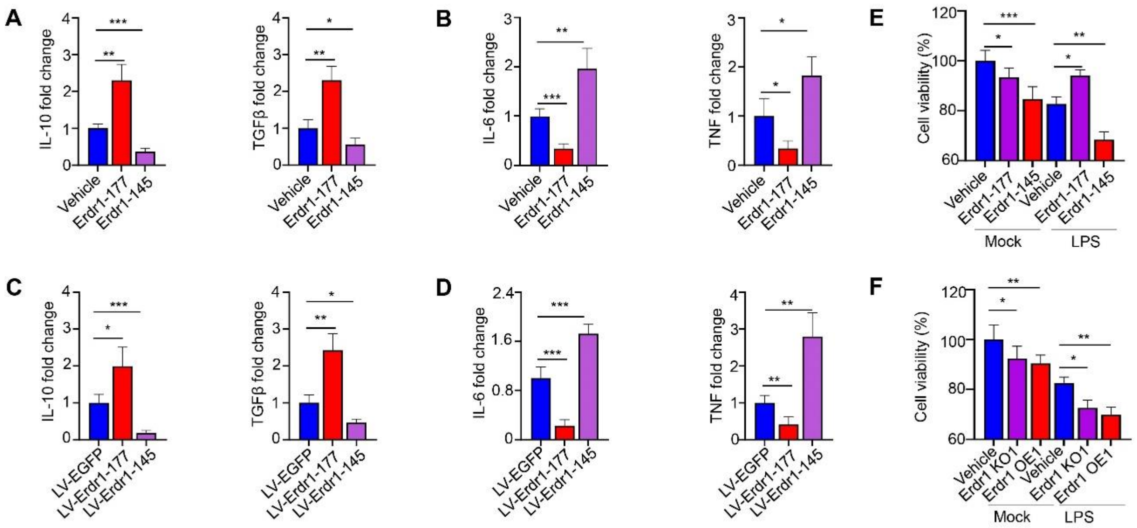


**Supplemental Figure 2 Erdr1 mediates macrophage polarization and modulates cell death**

**(A-B)** BMDM were treated with Erdr1-177 protein (1 µg/ml) or Erdr1-145 (1 µg/ml) or vehicle (1 µg/ml BSA) for 48 hours and samples were collected for the following analysis: qPCR analysis of M1 marker gene IL-10 and TGFβ **(A)** and M2 marker gene IL-6 and TNF **(B)** (n=4). **(C-D)** BMDM were transfected lentivirus LV-Erdr1-177 or LV-Erdr1-145 or control (LV-EGFP) for 48 hours and samples were collected for the following analysis: qPCR analysis of M1 marker gene IL-10 and TGFβ **(C)** and M2 marker gene IL-6 and TNF **(D)** (n=4). qPCR normalized to β-actin expression. **(E)** BMDM were treated as follows for 48 hours: 1. Vehicle; 2. Erdr1-177 (1µg/ml) 3. Erdr1-145 (1µg/ml); 4. Vehicle+ LPS (1µg/ml); 5. Erdr1-177 (1µg/ml) + LPS (1µg/ml); 6. Erdr1-145 (1µg/ml) + LPS (1µg/ml), MTT assay was performed for measuring cell viability (n=6). **(F)** BMDM were genetic knockout of Erdr1 (Erdr1 KO) or overexpressed Erdr1 (Erdr1 OE) by lentivirus transduction for 40 hours, the vehicle represents transduction with vehicle control lentivirus. Cells were treated without or with 1µg/ml LPS treatment for 48 hours. MTT assay was performed for measuring cell viability (n=6). Data represent mean ± SD. $*p<0.05$, $**p<0.01$, $***p<0.001$.
